# Legacy of draught cattle breeds of South India: Insights into population structure, genetic admixture and maternal origin

**DOI:** 10.1101/2021.01.21.427560

**Authors:** Vandana Manomohan, Ramasamy Saravanan, Rudolf Pichler, Nagarajan Murali, Karuppusamy Sivakumar, Krovvidi Sudhakar, Raja K Nachiappan, Kathiravan Periasamy

**Affiliations:** Animal Production and Health Section, Joint FAO/IAEA Division of Nuclear Techniques in Food and Agriculture, International Atomic Energy Agency, Vienna, Austria; Veterinary College and Research Institute, Namakkal, Tamil Nadu Veterinary and Animal Sciences University, Chennai, India; NTR College of Veterinary Science, Gannavaram, Sri Venkateswara Veterinary University, Tirupati, Andhra Pradesh, India; National Bureau of Animal Genetic Resources, Karnal, Haryana, India; Animal Genetics Resources Branch, Animal Production and Health Division, Food and Agriculture Organization of the United Nations, Rome, Italy

**Keywords:** Zebu, diversity, inbreeding, genetic relationship, taurine introgression, haplogroup, mismatch distribution, domestication

## Abstract

The present study is the first comprehensive report on diversity, population structure, genetic admixture and mitochondrial DNA variation in South Indian draught type zebu cattle. The diversity of South Indian cattle was moderately higher. A significantly strong negative correlation coefficient of −0.674 (P<0.05) was observed between the effective population size of different breeds and their estimated F_IS_. The phylogeny and genetic structure analysis revealed the distinctness of Kangayam, Vechur and Punganur cattle from the rest of the zebu breeds. The results showed the influence of Hallikar breed in the development of most Mysore type cattle breeds of South India with the exception of Kangayam. Bayesian clustering analysis was performed to assess the taurine admixture in South Indian zebu cattle using purebred Jersey and Holstein-Friesian as reference genotypes. Relatively high levels of taurine admixture (>6.25%) was observed in Punganur, Vechur, Umblachery and Pulikulam cattle breeds. Two major maternal haplogroups, I1 and I2, typical of zebu cattle were observed, with the former being predominant than the later. The pairwise differences among the I2 haplotypes of South Indian cattle were relatively higher than West Indian (Indus valley site) zebu cattle. The results indicated the need for additional sampling and comprehensive analysis of mtDNA control region variations to unravel the probable location of origin and domestication of I2 zebu lineage. The present study also revealed major concerns on South Indian zebu cattle (i) risk of endangerment due to small effective population size and high rate of inbreeding (ii) lack of sufficient purebred zebu bulls for breeding and (iii) increasing level of taurine admixture in zebu cattle. Availability of purebred semen for artificial insemination, incorporation of genomic/molecular information to identify purebred animals and increased awareness among farmers will help to maintain breed purity, conserve and improve these important draught cattle germplasms of South India.

## 1. INTRODUCTION

Ever since domestication, animals have been used as an important source of draught power for agricultural production. Among them, cattle have been used for a variety of farm operations like ploughing, carting, tilling, sowing, weeding, water lifting, threshing, oil extraction, sugarcane crushing and transport. Over time, with increasing mechanization of agriculture, the use of draught cattle in farm operations reduced significantly and became negligible in advanced economies. However, in developing countries, particularly in small holder production systems and in hilly and mountainous regions, agricultural production still relies on draught animal power (Zhou et al. 2018). For example, 55% of the smallholder farmers in Swaziland rely on draft animal power for land cultivation, and more than 88% of draft animals found on the Swazi National Land are cattle (Kienzle et al. 2013). In India, the significance of draught animal power has reduced greatly in the last few decades with its share in total farm power availability declining from about 78% in 1960-61 to 7% in 2013-2014 (Singh et al. 2014) and expected to further reduce to around 5% by 2020 (Natarajan et al. 2016). Over the years, the annual use of draught animals declined from about 1200-1800 hours to 300-500 hours per pair per year. Consequently, the population of draught cattle steadily decreased at a negative annual compound growth rate of about −0.8%, resulting in significant decline of purebred native breeds.

India is home to the largest cattle population (13.1 % of world’s cattle population) in the world which constitutes 37.3 % of its total livestock. As per the 19th livestock census, India has 190.9 million cattle, out of which 151 million is indigenous cattle. The diversity of Indian cattle is represented by 60 local breeds, eight regional transboundary breeds and seven international transboundary breeds (FAO, 2015). Of these, 41 have been registered by National Bureau of Animal Genetic Resources, the nodal agency for the registration of livestock breeds in India. The Indian Zebu cattle (*Bos indicus*) breeds have been classified into three major groups: dairy type, draught type and dual-purpose cattle. The dairy type and dual-purpose cattle breeds are predominantly distributed in North and North-Western India, while most indigenous draught type cattle breeds are located in Southern and Eastern India. Apart from rapid mechanization of agriculture in South India, the increasing cost of maintaining draught type cattle forced the farmers to replace these animals with crossbred cattle that fetched better returns from the sale of milk to dairy cooperatives. During late 1980s and early 1990s, the draught cattle breeds started losing their economic relevance and the farmers increasingly resorted to crossbreeding with commercial taurine cattle like Jersey, Holstein-Friesian and Brown Swiss. Such crossbred cows approximately yielded more than double the milk as that of indigenous draught breeds. Combined with farmers’ preference for crossbreds, the relatively easy access to breeding services (through state animal husbandry departments, state cooperatives and private inseminators, etc.) and availability of better infrastructure for artificial insemination and veterinary care, the proportion of crossbred cattle among the breedable females became much higher in South Indian states than rest of India. The crossbred cattle constitute 35.9% of total cattle in Andhra Pradesh, 41.9% in Karnataka, 76.1% in Tamil Nadu and 94.6% in Kerala as against the national average of 27.7%. This resulted in significant genetic dilution of draught cattle breeds of South India with varying levels of zebu-taurine admixture.

Indiscriminate crossbreeding, lack of stabilization of exotic inheritance and absence of selection among F2 hybrids led to negative impacts like increased disease susceptibility, decreased fertility and drop in production levels among crossbred cattle. During the last two decades, there has been a renewed interest in the conservation of purebred indigenous draught cattle breeds and implementation of selective breeding programs targeting a range of traits. Selective breeding schemes among certain purebred native breeds like Kangayam, Ongole, Deoni and Hallikar were put in place by making available the purebred frozen semen for artificial insemination. Such programs have been constrained by the availability of superior genetic merit bulls and unavailability of sufficient of number of true to the type, purebred bulls. With the absence of pedigree records, the level of taurine inheritance in most purebred draught cattle populations is not clearly understood. Further, information on genetic differentiation, population structure and maternal inheritance among South Indian cattle breeds is scanty. Hence, the present study was undertaken with the following objectives: (i) assess biodiversity and population structure among purebred South Indian draught cattle breeds (ii) estimate level of genetic admixture among purebred zebu cattle and (iii) identify mitochondrial DNA based maternal lineages and evaluate phylogeography of South Indian cattle.

## 2. MATERIALS AND METHODS

### 2.1. Animal ethics statement

Blood samples were collected by jugular venipuncture into EDTA vacutainer tubes from various purebred zebu, commercial taurine and crossbred cattle. Blood sampling was performed by local veterinarians in the respective native breed tracts following the standard good animal practice. The consent was obtained from animal owners for the usage of the samples in this study. Therefore, no further license from the “Institutional Committee for Care and Use of Experimental Animals” of the Tamil Nadu Veterinary and Animal Sciences University, Chennai, India and the Joint FAO/IAEA Division, International Atomic Energy Agency, Vienna, Austria was required.

### 2.2. Sample collection

A total of 529 blood samples were collected from cattle representing 14 breeds/populations that are located in South/Peninsular Indian states: Ongole (n=51) and Punganur (n=18) from Andhra Pradesh; Hallikar (n=36) and Deoni (n=53) from Karnataka; Vechur (n=26) from Kerala; Alambadi (n=29), Bargur (n=52), Kangayam (n=51), Pulikulam (n=34), Umblachery (n=33), Holstein-Friesian (n=15), Jersey (n=34), Holstein-Friesian crossbred (n=39) and Jersey crossbred (n=58) from Tamil Nadu. A stratified random sampling procedure was followed to collect blood from unrelated cattle with typical phenotypic features and located in randomly selected villages of the native breed tract. With the absence of pedigree records in most instances, unrelatedness was ensured by interviewing the farmers about the breeding history. Genomic DNA was extracted by standard phenol chloroform method (Sambrook and Russell, 2001). The DNA samples were subjected to quality control using Nanodrop spectrophotometer and stored at −20°C until further processing.

### 2.3. Microsatellite genotyping

All the DNA samples were subjected to PCR amplification using 27 FAO recommended microsatellite markers (FAO, 2011; Supplementary Table ST1). The forward primers conjugated to one of the three fluorescent dyes (FAM, HEX and ATTO550) were used for genotyping these marker loci. Polymerase chain reaction was performed under following conditions: initial denaturation for 5 min at 95°C, followed by 35 cycles of denaturation for 30s at 95°C, 1 minute at specific annealing temperatures of marker locus, elongation at 72°C for 1 min with the final extension at 72°C for 10 min. The PCR products were then electrophoresed after multiplexing in an automated DNA analyzer ABI3100 (Applied Biosystems, USA) with ROX500 (Applied Biosystems, USA) as internal lane control. All the 27 STR loci were multiplexed in six sets for genotyping as shown in Supplementary Table ST1. Determination of allele size and extraction of genotype data was performed using GENEMAPPER software.

### 2.4. Sequencing mitochondrial DNA control region

Primers were custom designed and synthesized to amplify 1207 bp bovine mitochondrial DNA control region using Primer 3 version 4.0 (http://bioinfo.ut.ee/primer3-0.4.0/). The amplified mtDNA control region included nucleotide positions 15167 to 30 of the complete bovine mitochondrial genome sequence available at NCBI GenBank accession number AF492350 (Hiendleder et al. 2008). The location and sequence of forward and reverse primers are as follows: BTMTD1-F (Positions 15167-15186; 5’ AGGACAACCAGTCGAACACC 3’) and BTMTD1-R 5’ (Positions 11-30; GTGCCTTGCTTTGGGTTAAG 3’). The amplicon included complete D-loop (control region) flanked by partial cytochrome B, complete t-RNA-Thr and complete t-RNA-Pro coding sequences at 5’ end while partial t-RNA-Phe sequence flanked the 3’end. Polymerase chain reaction was performed in a total reaction volume of 40µl with the following cycling conditions: initial denaturation at 95°C for 15 min followed by 30 cycles of 95°C for 1 min; 58°C for 1 min; 72°C for 1 min with final extension at 72°C for 10 min. Purified PCR products were sequenced using Big Dye Terminator Cycle Sequencing Kit (Applied Biosystems, U.S.A) on an automated Genetic Analyzer ABI 3100 (Applied Biosystems, U.S.A).

### 2.5. Statistical analysis

The presence of null alleles in the dataset was checked using Microchecker version 2.2.3 (Oosterhout et al. 2004). The neutrality of microsatellite markers used in the present study was evaluated by F_ST_ outlier approach as implemented in LOSITAN version 1 for windows (Antao et al. 2008). Basic diversity indices (allelic diversity and heterozygosity) and genetic distances (pairwise Nei’s genetic distance and pairwise inter-individual allele sharing distance) among South Indian cattle were estimated using MICROSATELLITE ANALYZER (MSA) version 4.05 (Dieringer and Schlotterer, 2003). A UPGMA tree was constructed based on pairwise Nei’s genetic distances using PHYLIP version 3.5 (Felsenstein, 1993). F statistics for each locus (Weir and Cockerman, 1984) was calculated and tested using the program FSTAT version 2.9.3.2. The exact tests for Hardy-Weinberg Equilibrium to evaluate both heterozygosity deficit and excess at each marker locus in each population (HWE) were performed using GENEPOP software. Analysis of molecular variance (AMOVA) was performed using ARLEQUIN version 3.1 (Excoffier et al. 2005). Multidimensional scaling display of pairwise F_ST_ was conducted using SPSS version 13.0. Bayesian clustering analysis was performed to evaluate population structure and genetic admixture among South Indian cattle using STRUCTURE 2.3.4 (Pritchard et al. 2000). The program was run with the assumption of K=2 to K=15 with at least 20 runs for each K. The number of burn in periods and MCMC repeats used for these runs were 200000 and 200000 respectively. The procedure of Evanno et al. (2005) was used to estimate the second order rate of change in LnP(D) and identify the optimal K.

Mitochondrial DNA sequence editing, assembly of contigs and multiple sequence alignments were performed using Codon Code Aligner version 3.7.1. Mitochondrial DNA diversity parameters including nucleotide diversity, haplotype diversity, parsimony informative sites and average number of nucleotide differences were calculated using DnaSP, version 4.10 (Rozas, 2009). Frequency of different haplotypes, haplotype sharing and pairwise F_ST_ were estimated using ARLEQUIN 2.1. MEGA version 6.0 (Tamura et al. 2011) was utilized to identify optimal DNA substitution model and construct maximum-likelihood phylogeny. HKY+I+G was identified as the optimal model for the current sequence dataset based on Akaike information criterion (AIC). The following sequences were used as references for different maternal lineages reported in zebu and taurine cattle: I1 (AB085923, AB268563, AB268579, EF417970, EF417971 and L27733), I2 (AB085921, AB268559, AB268570, AB268573, AB268581 and AY378133), T1 (DQ124399, EU177841 and EU177842), T2 (DQ124383, DQ124393, EU177849 and EU177851), T3 (EU177815, EU177816, EU177838 and EU177839), T4 (DQ124372, DQ124375, DQ124377 and DQ124392), and T5 (EU177862, EU177863, EU177864 and EU177865). Additionally, 298 mtDNA control region sequences deposited to GenBank from Bhutan (n=121), Nepal (n=14), China (n=47), North India (n=53), West India (n=45) and Vietnam (n=18) were utilized for comparative analysis. The accession number and details of GenBank sequences used in the study are provided in Supplementary Table ST2. Pairwise F_ST_ derived from mtDNA haplotype frequency were utilized to perform principal components analysis using SPSS version 13.0. Mismatch distribution and the tests for demographic expansion were performed using ARLEQUIN 3.1. Three parameters viz. tau, theta 0 and theta 1 were estimated under the sudden expansion model and the tests for goodness of fit were performed based on sum of square deviations (SSD) between the observed and expected mismatch and Harpending’s raggedness index. The South Indian cattle breeds were subjected to tests for selective neutrality through estimation of Tajima’s D and Fu’s F_S_. Median Joining (MJ) network of South Indian cattle haplotypes was reconstructed using NETWORK 4.5.1.2 (Bandelt et al. 1999).

## 3. RESULTS

### 3.1. Basic diversity measures and Hardy-Weinberg equilibrium

In the present study, a total of 13932 genotypes were generated across 27 microsatellite loci in 516 cattle belonging to 10 draught type zebu breeds, 2 commercial taurine breeds and 2 crossbred populations. The mean basic diversity indices of different South Indian cattle breeds are presented in Table 1. Allelic diversity in terms of mean observed number of alleles varied from 5.93 (Punganur) to 8.07 (Pulikulam) across different zebu cattle breeds, while it was 4.7 and 5.81 in the commercial taurine breeds, Jersey and Holstein-Friesian respectively. The observed heterozygosity ranged from 0.598 (Kangayam) to 0.671 (Alambadi and Punganur) while the expected heterozygosity varied between 0.637 (Kangayam) and 0.727 (Punganur) among the South Indian zebu cattle. Similarly, allelic richness was lowest in Kangayam and highest in Pulikulam cattle. Assessment of within breed diversity based on inter-individual allele sharing distance revealed Kangayam with the lowest mean distance among individuals (0.568) while Pulikulam cattle showed the highest mean distance (0.647). The allelic diversity observed among South Indian zebu cattle was relatively low as compared to that of West-Central Indian (Shah et al. 2012), North Indian (Sharma et al. 2015) and East Indian (Sharma et al. 2013) cattle. The primary domestication centre of zebu (*Bos indicus*) cattle in North-Western part of India near Indus valley could explain the higher diversity of North Indian cattle compared to that of South Indian breeds. Despite the relatively low allelic diversity, the observed and expected heterozygosity of South Indian cattle were moderately higher (>60%) and comparable to that of West-Central, North and East Indian cattle (Shah et al. 2012; Sharma et al. 2013, 2015). As expected, the crossbred cattle (Jersey crossbreds and Holstein-Friesian crossbreds) exhibited a higher level of allelic diversity and heterozygosity than indigenous zebu cattle.

**Table 1.**
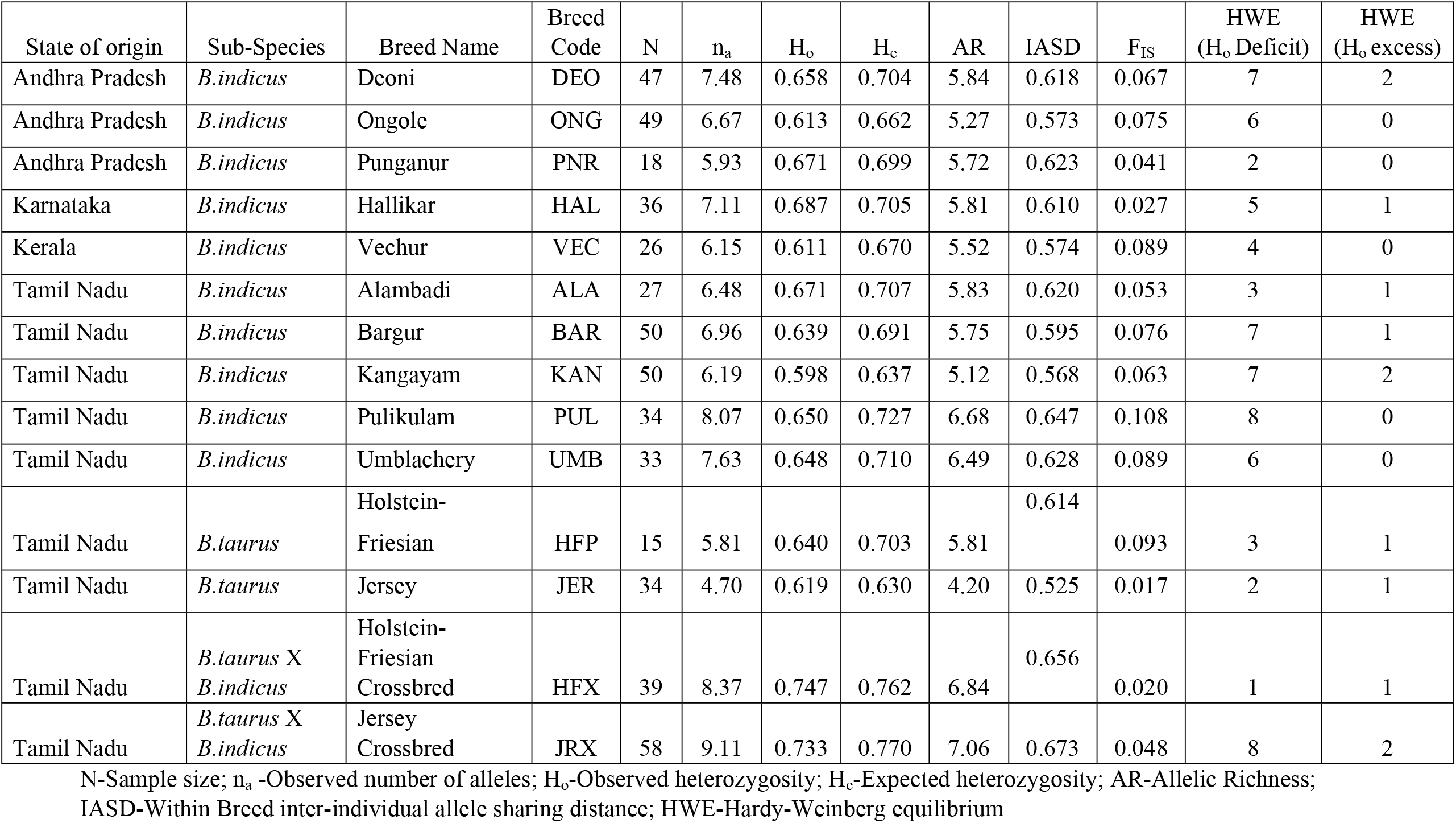
Summary diversity statistics for draught type zebu, taurine and crossbred cattle of South India

| State of origin | Sub-Species | Breed Name | Breed Code | N | n<sub>a</sub> | H<sub>o</sub> | H<sub>e</sub> | AR | IASD | F<sub>IS</sub> | HWE (H<sub>o</sub> Deficit) | HWE (H<sub>o</sub> excess) |
| --- | --- | --- | --- | --- | --- | --- | --- | --- | --- | --- | --- | --- |
| Andhra Pradesh | *B.indicus* | Deoni | DEO | 47 | 7.48 | 0.658 | 0.704 | 5.84 | 0.618 | 0.067 | 7 | 2 |
| Andhra Pradesh | *B.indicus* | Ongole | ONG | 49 | 6.67 | 0.613 | 0.662 | 5.27 | 0.573 | 0.075 | 6 | 0 |
| Andhra Pradesh | *B.indicus* | Punganur | PNR | 18 | 5.93 | 0.671 | 0.699 | 5.72 | 0.623 | 0.041 | 2 | 0 |
| Karnataka | *B.indicus* | Hallikar | HAL | 36 | 7.11 | 0.687 | 0.705 | 5.81 | 0.610 | 0.027 | 5 | 1 |
| Kerala | *B.indicus* | Vechur | VEC | 26 | 6.15 | 0.611 | 0.670 | 5.52 | 0.574 | 0.089 | 4 | 0 |
| Tamil Nadu | *B.indicus* | Alambadi | ALA | 27 | 6.48 | 0.671 | 0.707 | 5.83 | 0.620 | 0.053 | 3 | 1 |
| Tamil Nadu | *B.indicus* | Bargur | BAR | 50 | 6.96 | 0.639 | 0.691 | 5.75 | 0.595 | 0.076 | 7 | 1 |
| Tamil Nadu | *B.indicus* | Kangayam | KAN | 50 | 6.19 | 0.598 | 0.637 | 5.12 | 0.568 | 0.063 | 7 | 2 |
| Tamil Nadu | *B.indicus* | Pulikulam | PUL | 34 | 8.07 | 0.650 | 0.727 | 6.68 | 0.647 | 0.108 | 8 | 0 |
| Tamil Nadu | *B.indicus* | Umblachery | UMB | 33 | 7.63 | 0.648 | 0.710 | 6.49 | 0.628 | 0.089 | 6 | 0 |
| Tamil Nadu | *B.taurus* | Holstein-Friesian | HFP | 15 | 5.81 | 0.640 | 0.703 | 5.81 | 0.614 | 0.093 | 3 | 1 |
| Tamil Nadu | *B.taurus* | Jersey | JER | 34 | 4.70 | 0.619 | 0.630 | 4.20 | 0.525 | 0.017 | 2 | 1 |
| Tamil Nadu | *B.taurus* X *B.indicus* | Holstein-Friesian Crossbred | HFX | 39 | 8.37 | 0.747 | 0.762 | 6.84 | 0.656 | 0.020 | 1 | 1 |
| Tamil Nadu | *B.taurus* X *B.indicus* | Jersey Crossbred | JRX | 58 | 9.11 | 0.733 | 0.770 | 7.06 | 0.673 | 0.048 | 8 | 2 |

N-Sample size; n<sub>a</sub> -Observed number of alleles; H<sub>o</sub>-Observed heterozygosity; H<sub>e</sub>-Expected heterozygosity; AR-Allelic Richness; IASD-Within Breed inter-individual allele sharing distance; HWE-Hardy-Weinberg equilibrium

The mean estimated F_IS_ values were positive in all the zebu cattle breeds and ranged between 0.027 (Hallikar) and 0.108 (Pulikulam). Breeds like Umblachery, Vechur, Ongole and Bargur showed mean F_IS_ greater than 7% indicating significant heterozygosity deficit, possibly due to close breeding and potential inbreeding. The mean F_IS_ estimates observed in most South Indian cattle breeds were higher than those reported for East Indian (Sharma et al. 2013) and most North Indian cattle with the exception of Mewati and Gaolao (Sharma et al. 2015). However, the mean F_IS_ estimates reported for West-Central Indian cattle were much higher than that of South Indian cattle (Shah et al. 2012). Similarly, the mean F_IS_ for Vechur and Ongole cattle observed in the present study was significantly lower than reported previously by Radhika et al. (2018) and Sharma et al. (2015) respectively. The test for Hardy-Weinberg equilibrium (HWE) was conducted on a total of 378 breed X locus combinations. Among the 270 breed X locus combinations tested for zebu cattle, about 62 (22.96%) deviated significantly (P<0.05) from HWE, of which 55 (20.37%) were due to heterozygosity deficit and 7 (2.59%) were due to heterozygosity excess. The number of loci deviating from HWE varied from two (Punganur) to nine (Deoni and Kangayam) across the investigated cattle breeds. Heterozygosity deficiency could be due to several reasons that may include population sub-division (Wahlund effect), null alleles, selective forces operating at certain loci, scoring bias, small sample size and inbreeding. Microchecker analysis did not reveal the presence of any significant null allele in the dataset.

To evaluate the influence of selection on the loci under study, a F_ST_ outlier approach was followed. Loci under the influence of positive/directional selection are expected to show relatively higher F_ST_ values (outliers) as compared to the loci following neutral genetic variation. The loci under balancing selection are expected to show relatively lower F_ST_ values (outliers) (Excoffier et al. 2009; Vitti et al. 2013). This selection detection strategy evaluates the relationship between F_ST_ and expected heterozygosity under an island model of migration and describes an expected distribution of F_ST_ vs H_e_ under selective neutrality. This distribution is utilized to identify the outlier loci that have excessively high or low F_ST_ as compared to neutral expectations. Loci outside the 99% confidence interval and false discovery rate less than 1% were identified as candidates affected by positive and balancing selection (Antao et al. 2008). The results revealed two loci, HEL5 (F_ST_=0.215) and HEL13 (F_ST_=0.216) as significant outliers with high F_ST_ values, indicating positive selection in force (Supplementary Figure SF1). Of these, HEL5 exhibited significant heterozygosity deficit and consequent deviation from HWE in 12 out of 14 studied populations. Both the loci have been reported to be associated with production and/or reproduction traits in cattle (de Oliveira et al. 2005; Li et al. 2010; Vanessa et al. 2018). Specifically, the homozygous long alleles at HEL5 (allele size 159-169) were significantly associated (p=0.022) with long calving interval in Brangus-Ibage cattle, a composite breed with taurine and zebu ancestry (de Oliveira et al. 2005). In the present study, significant difference in the frequency of long (allele size 160-180) and short alleles (allele size 148-158) was observed among the zebu, taurine and crossbred cattle, resulting in higher estimation of F_ST_. Selective processes affect neutral loci when the latter is in linkage disequilibrium with other loci subjected to selection. Considering the possibility of selective forces operating at the two F_ST_ outlier loci, HEL5 and HEL13 were removed from subsequent analysis of genetic relationship, admixture and population structure of South Indian cattle.

### 3.2. Genetic differentiation and phylogeny

The global F statistics among the South Indian cattle breeds are presented in Supplementary Table ST3. The overall global F_IT_, F_ST_ and F_IS_ among zebu, taurine and crossbred cattle was 0.143, 0.101 and 0.047 respectively, while those estimates among zebu cattle alone were 0.109, 0.057 and 0.056 respectively. The results indicated 10.1% of the total genetic variation among all the studied cattle were due to between breed differences and it reduced to 5.7% when only zebu cattle breeds were considered. The global F_ST_ among all cattle breeds/populations ranged from 0.052 (INRA005) to 0.223 (ETH152), whereas among zebu cattle breeds, it varied between 0.024 (INRA005) and 0.114 (HAUT24). Among the investigated loci, ETH152 provided relatively more information on zebu-taurine divide while HAUT24 provided significant information on differences among zebu cattle breeds. The global F_ST_ among South Indian zebu cattle observed in the present study was comparable to that of East Indian zebu (Sharma et al. 2013) cattle. However, much higher genetic differentiation had been reported among West-Central (Shah et al. 2012) and North Indian (Sharma et al. 2015) cattle breeds. The pairwise F_ST_ and Nei’s genetic distance among the draught type zebu, taurine and crossbred cattle are presented in Table 2. The pairwise F_ST_ ranged from 0.0234 (Hallikar-Alambadi) to 0.107 (Kangayam-Punganur) among the South Indian zebu cattle breeds. Alambadi cattle was observed to be genetically less differentiated from Hallikar (F_ST_=0.0234), Pulikulam (F_ST_=0.0237) and Bargur (F_ST_=0.0259) while Kangayam cattle showed highest differentiation from Punganur (F_ST_=0.107), Vechur (F_ST_=0.1034) and Hallikar (F_ST_=0.0933). Interestingly, significantly high genetic differentiation (F_ST_=0.0919) was observed among the two dwarf/miniature type Vechur and Punganur cattle. Similar findings were observed with respect to the estimates of pairwise Nei’s genetic distance among the South Indian zebu cattle breeds. The radial tree and dendrogram constructed following UPGMA algorithm based on pairwise Nei’s genetic distance is presented in Figure 1. Broadly, the taurine, zebu and their crossbred cattle clustered separately as expected. Among the zebu cattle, a major cluster was formed by Alambadi, Bargur, Umblachery, Pulikulam, Deoni, Ongole and Hallikar, within which three sub-clusters were observed. Alambadi and Bargur formed the first sub-cluster, Umblachery and Pulikulam formed the second sub-cluster while Deoni, Ongole and Hallikar formed the third sub-cluster. Kangayam, Vechur and Punganur cattle breeds did not form part of any of these three clusters and were distinct among themselves.

**Table 2.** Pairwise F_ST_ (upper triangle) and pairwise Nei’s genetic distance (lower triangle) among draught type zebu, taurine and crossbred cattle from South India

|  | ALA | BAR | DEO | HAL | HFP | HFX | JER | JRX | KAN | ONG | PNR | PUL | UMB | VEC |
| --- | --- | --- | --- | --- | --- | --- | --- | --- | --- | --- | --- | --- | --- | --- |
| ALA | - | 0.026 | 0.037 | 0.023 | 0.172 | 0.115 | 0.218 | 0.092 | 0.063 | 0.053 | 0.048 | 0.024 | 0.037 | 0.060 |
| BAR | 0.0744 | - | 0.041 | 0.050 | 0.190 | 0.130 | 0.227 | 0.101 | 0.077 | 0.050 | 0.077 | 0.034 | 0.031 | 0.065 |
| DEO | 0.1154 | 0.0963 | - | 0.062 | 0.180 | 0.125 | 0.225 | 0.098 | 0.063 | 0.044 | 0.064 | 0.026 | 0.038 | 0.059 |
| HAL | 0.0882 | 0.1042 | 0.1487 | - | 0.173 | 0.116 | 0.212 | 0.096 | 0.093 | 0.079 | 0.059 | 0.043 | 0.045 | 0.073 |
| HFP | 0.4437 | 0.4347 | 0.4509 | 0.4321 | - | 0.007 | 0.094 | 0.052 | 0.222 | 0.203 | 0.175 | 0.174 | 0.168 | 0.187 |
| HFX | 0.2714 | 0.2627 | 0.2894 | 0.2594 | 0.0992 | - | 0.068 | 0.019 | 0.164 | 0.143 | 0.116 | 0.117 | 0.108 | 0.125 |
| JER | 0.4849 | 0.4730 | 0.5024 | 0.4582 | 0.2238 | 0.1749 | - | 0.045 | 0.262 | 0.257 | 0.216 | 0.215 | 0.206 | 0.246 |
| JRX | 0.2323 | 0.2138 | 0.2306 | 0.2250 | 0.1653 | 0.0621 | 0.1399 | - | 0.133 | 0.127 | 0.099 | 0.091 | 0.086 | 0.111 |
| KAN | 0.1321 | 0.1260 | 0.1326 | 0.1750 | 0.4815 | 0.3164 | 0.5241 | 0.2623 | - | 0.075 | 0.107 | 0.070 | 0.070 | 0.103 |
| ONG | 0.1221 | 0.1017 | 0.1078 | 0.1537 | 0.4625 | 0.3010 | 0.5344 | 0.2625 | 0.1310 | - | 0.077 | 0.054 | 0.050 | 0.070 |
| PNR | 0.1563 | 0.1760 | 0.1828 | 0.1611 | 0.4232 | 0.2725 | 0.4550 | 0.2380 | 0.2171 | 0.1657 | - | 0.051 | 0.068 | 0.092 |
| PUL | 0.0973 | 0.0918 | 0.0959 | 0.1273 | 0.4084 | 0.2394 | 0.4488 | 0.1885 | 0.1284 | 0.1158 | 0.1624 | - | 0.034 | 0.056 |
| UMB | 0.1144 | 0.0932 | 0.1094 | 0.1221 | 0.3817 | 0.2187 | 0.4157 | 0.1802 | 0.1324 | 0.1026 | 0.1748 | 0.0912 | - | 0.058 |
| VEC | 0.1586 | 0.1545 | 0.1544 | 0.1672 | 0.4115 | 0.2637 | 0.4840 | 0.2378 | 0.2049 | 0.1508 | 0.2048 | 0.1460 | 0.1408 | - |

**Figure 1.**
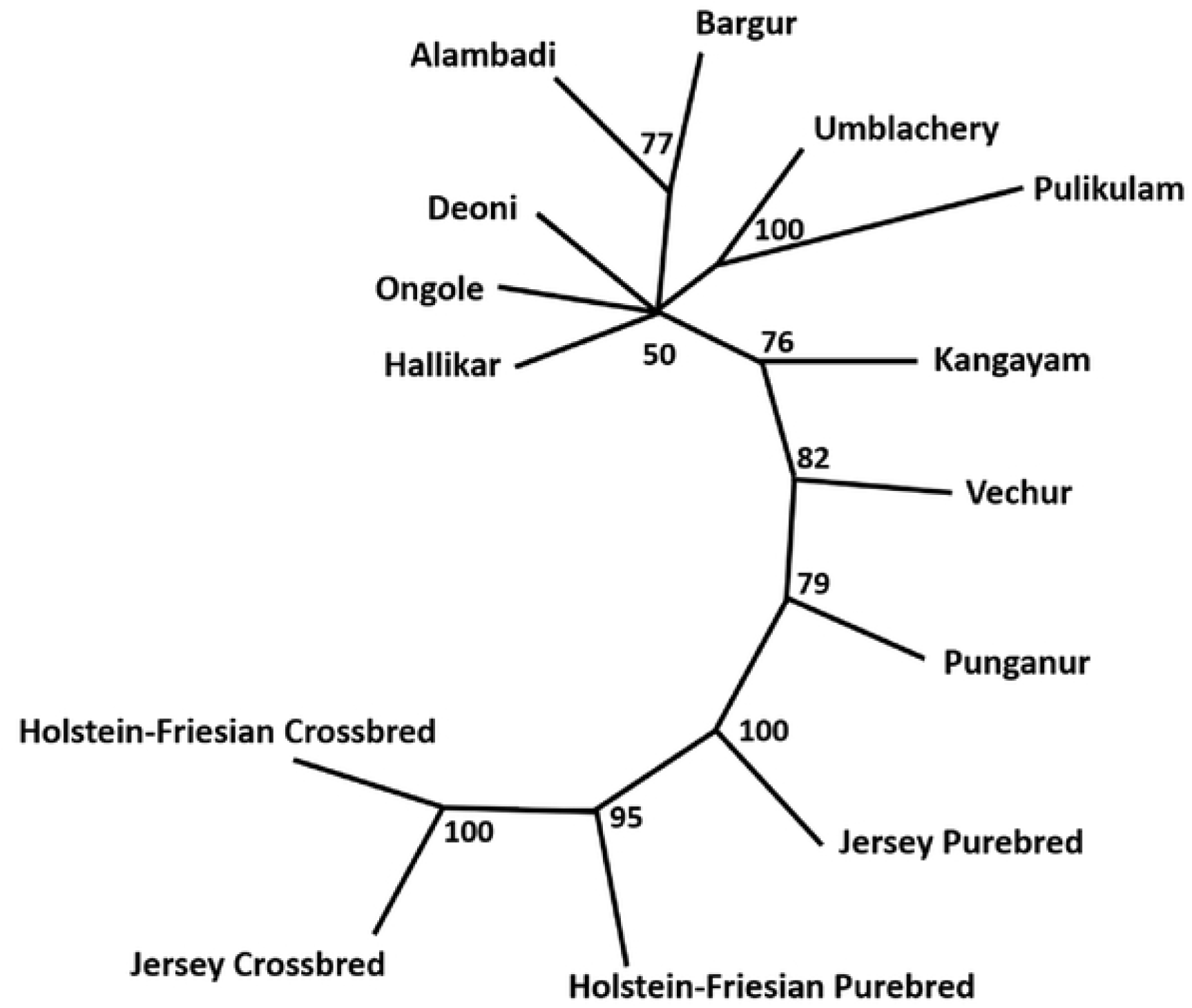
UPGMA tree derived from pairwise Nei’s genetic distance among zebu, taurine and crossbred cattle (numbers at nodes indicate percent bootstrap values of 10000 resampled data sets).

### 3.3. Genetic relationship and population structure

Understanding genetic relationship and detecting the cryptic genetic structure is an important first step to elucidate the demography and ancestral history of cattle populations. The multidimensional scaling (MDS) analysis were used to investigate the relationship among the studied cattle breeds. Multidimensional scaling (MDS) is an ordination technique used to display information contained in a distance matrix and to visualize relationship among variables within a dataset. This method helps to reduce the dimension of data and to detect structure among populations in a typical genetic diversity study. The MDS plots derived from pairwise F_ST_ values are presented in Figure 2. The observed s-stress value was 0.01202 when all the breeds/populations were considered while it was 0.08001 when the zebu cattle alone were considered. The MDS plots showed a pattern of clustering similar to that of phylogeny. The taurine and crossbreds clustered separately, while Kangayam, Vechur and Punganur cattle were distinct from rest of the zebu cattle breeds.

**Figure 2.**
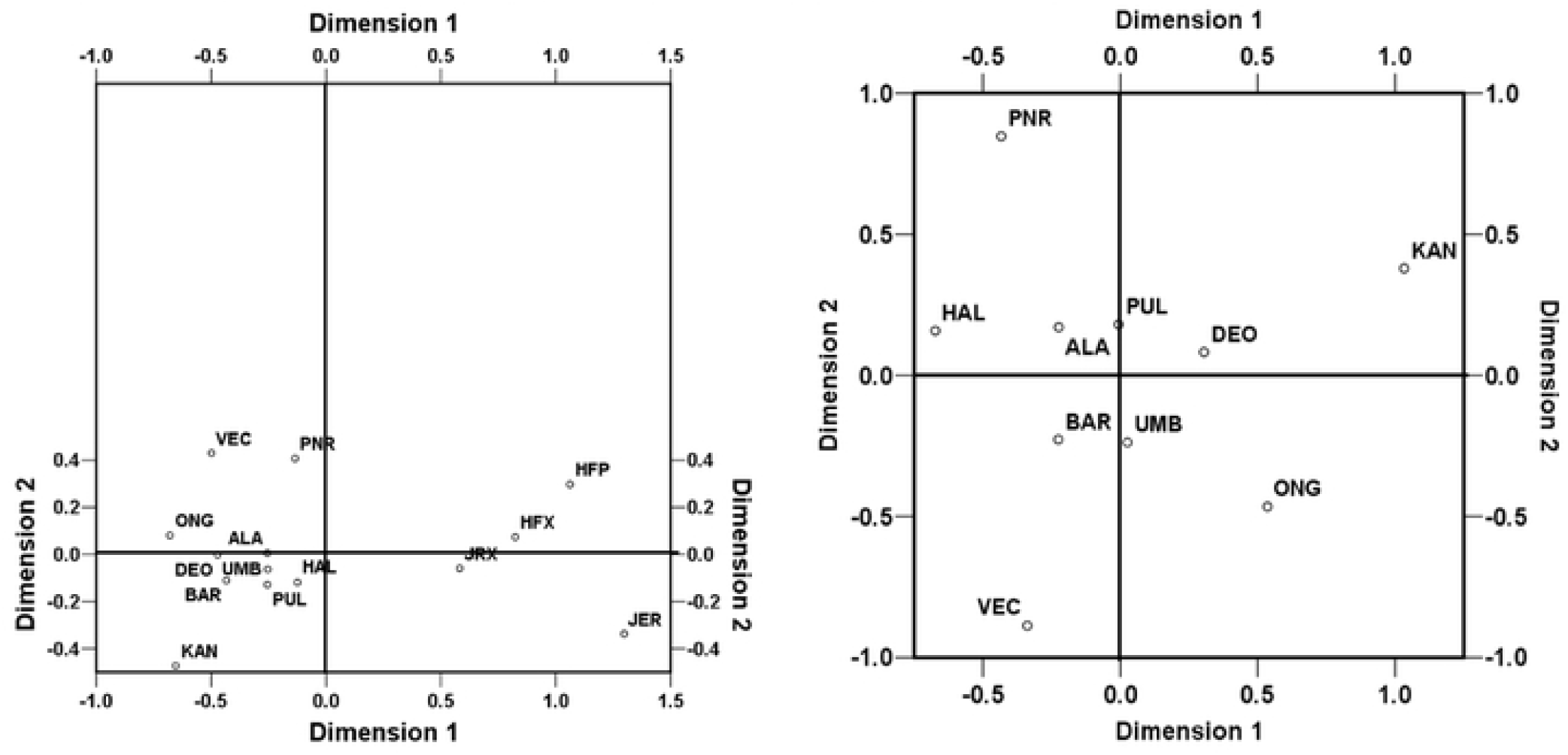
Multidimensional scaling display of pairwise F_ST_ among South Indian cattle (a) draught type zebu, taurine and crossbred cattle (S-Stress=0.01202) and (b) among draught type zebu cattle only (S-stress=0.08001)

Bayesian clustering of individuals without prior population information was performed to assess the cryptic genetic structure in the studied cattle breeds. Among K=2 to 15, the dataset was best described with K=2 genetic clusters at which the second order rate of change in LnP(D) was maximum (Supplementary Figure SF2). At K=2, the clustering of individuals followed the zebu-taurine divide (Figure 3). When K=3 was assumed, Zebu cattle breeds got clustered based on Hallikar or Kangayam ancestry. Both Hallikar and Kangayam cattle were assigned to their respective clusters with >90% proportion of membership coefficient. Deoni (80%) and Ongole (69.2%) were predominantly assigned to Kangayam ancestral cluster while Vechur (86.4%), Punganur (78.7%), Bargur (75.7%) and Alambadi (72.8%) cattle were predominantly assigned to Hallikar cluster. Pulikulam and Umblachery cattle showed admixture from both the ancestry. When K=4 was assumed, Deoni and Ongole clustered together independent of Kangayam ancestry while at K=5, Ongole clustered distinctly from that of Deoni. At K=6, Deoni and Bargur formed a separate cluster each, while Punganur and Vechur clustered together. At K=7, the miniature breeds, Pungaur and Vechur cattle clustered distinctly. Significant genetic admixture of zebu ancestry was observed among Alambadi, Pulikulam and Umblachery cattle consistently under the assumption of K=5 to 8.

**Figure 3.**
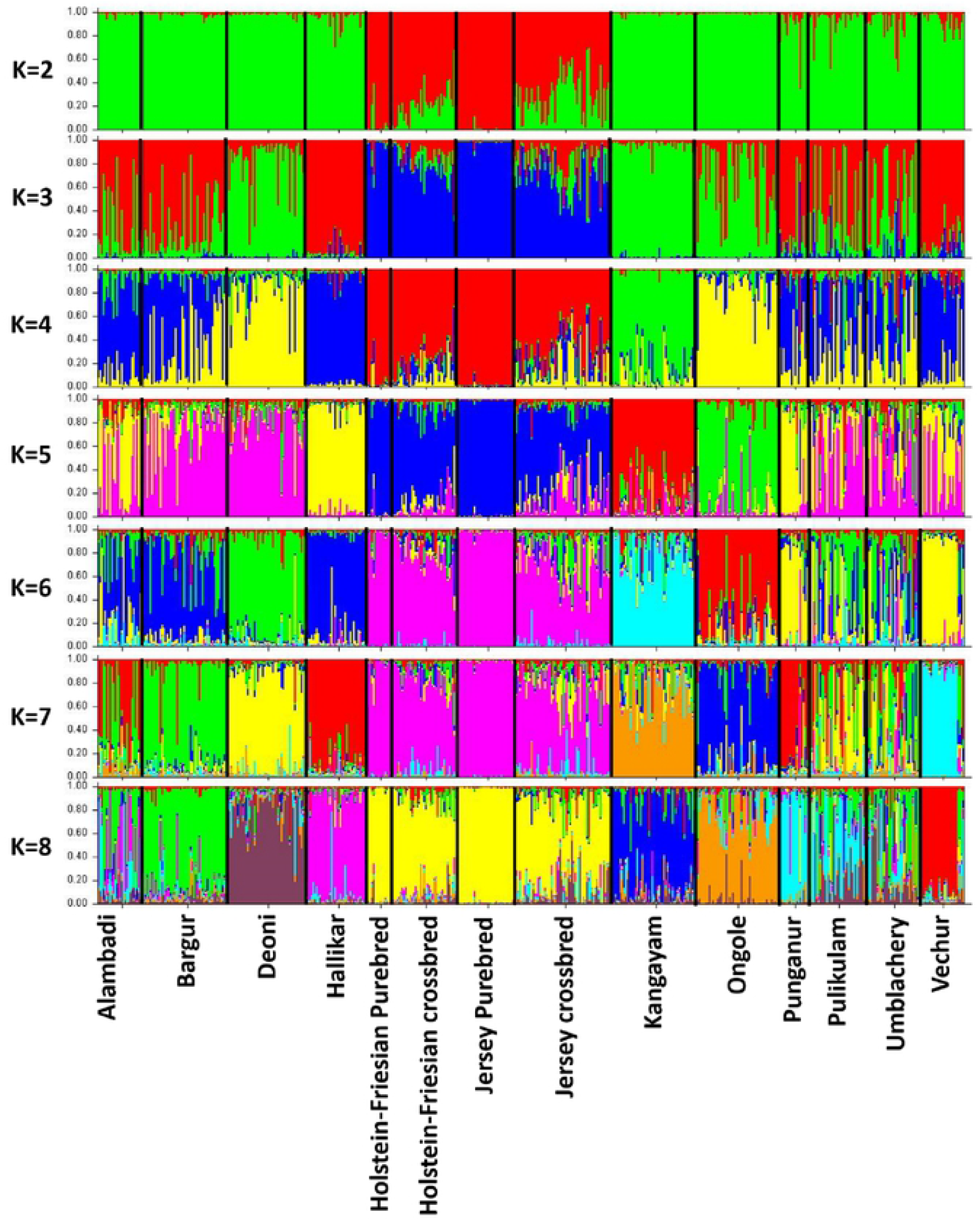
Bayesian clustering of 516 cattle under assumption of 2 to 8 clusters without a priori population information. The population names are given below the box plot with the individuals of different populations separated by vertical black lines

To further understand the degree of phylogeographic structure in the studied cattle breeds, analysis of molecular variance (AMOVA) was performed to assess the distribution of microsatellite variation as a function of both breed membership and geographic origin (Table 3). When no grouping was assumed, 94.56% of the total variance among zebu cattle was found within breeds while 5.44% of total variance was explained by between breed differences. When the breeds were grouped according to their province of origin (Grouping I – based on state/province), about 0.32% of total variance was due to differences among groups (P>0.05) while 5.21% was due to differences among populations within groups. When the grouping was done based on the classification of zebu cattle by Joshi and Phillips (1953) (Grouping II), among group variance was negligible (0.28%; P>0.05), while most of the between population variance was within groups (5.26%; P<0.05). When the grouping was done based on phylogeny (Grouping III), among group variance increased to 1.19%, but was not statistically significant (P>0.05) while variance among populations within groups was 4.41%. When the grouping was based on Bayesian genetic structure (Grouping IV; Group1-DEO; Group2-ONG; Group3-BAR, ALA, HAL, PUL, UMB; Group4-VEC; Group5-PNR; Group6-KAN), among group variance was 2.78% and significant (P<0.05) while variance among population within groups was 3.21%. The results of AMOVA thus indicated the structure based on Bayesian clustering to be evidently stronger than the putative structures based on provincial origin, Joshi and Phillips (1953) classification and phylogeny.

**Table 3.**
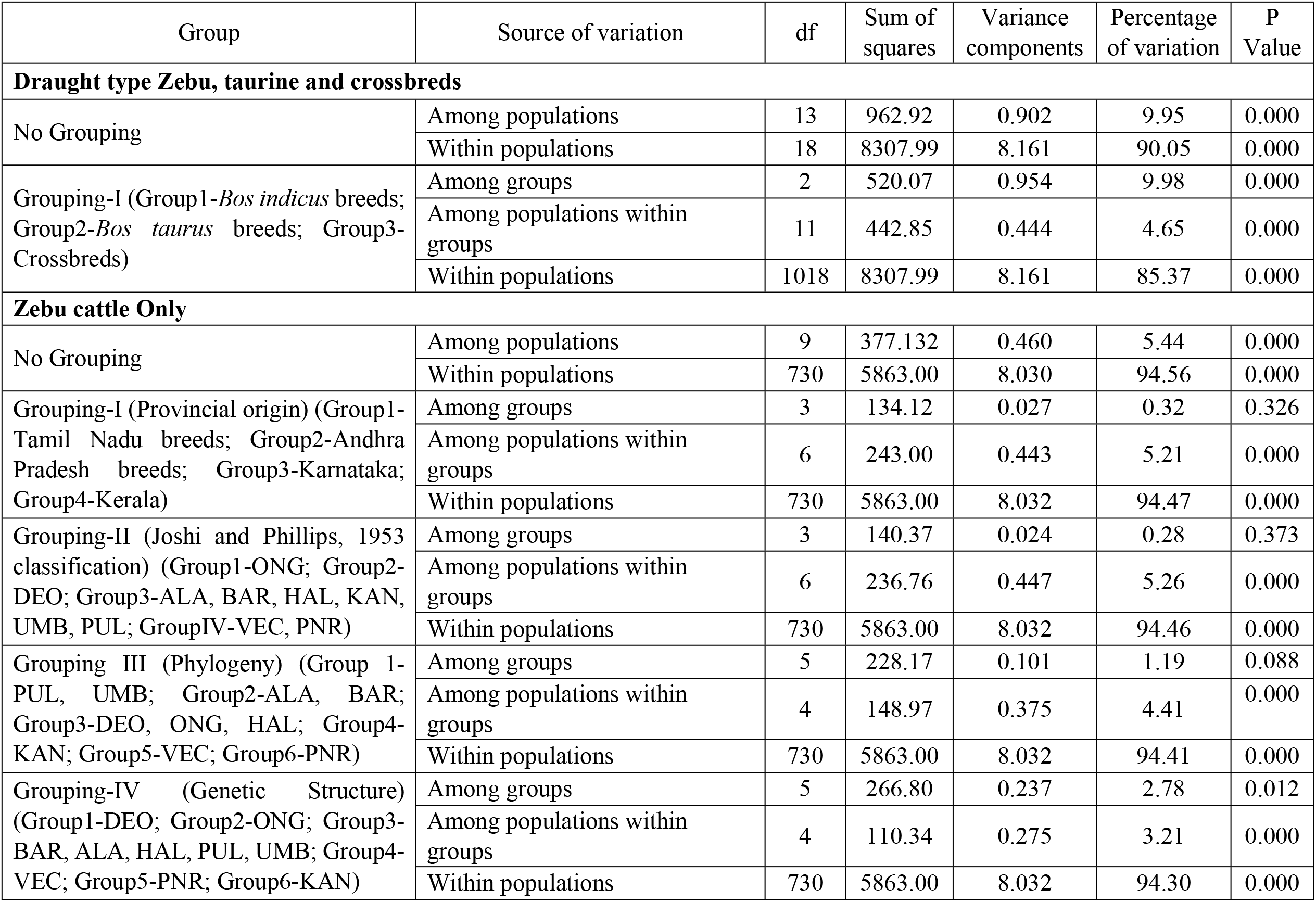
Analysis of variance (AMOVA) among draught type zebu, taurine and crossbred cattle from South India

| Group | Source of variation | df | Sum of squares | Variance components | Percentage of variation | P Value |
| --- | --- | --- | --- | --- | --- | --- |
| **Draught type Zebu, taurine and crossbreds** |  |  |  |  |  |  |
| No Grouping | Among populations | 13 | 962.92 | 0.902 | 9.95 | 0.000 |
|  | Within populations | 18 | 8307.99 | 8.161 | 90.05 | 0.000 |
| Grouping-I (Group1-*Bos indicus* breeds; Group2-*Bos taurus* breeds; Group3-Crossbreds) | Among groups | 2 | 520.07 | 0.954 | 9.98 | 0.000 |
|  | Among populations within groups | 11 | 442.85 | 0.444 | 4.65 | 0.000 |
|  | Within populations | 1018 | 8307.99 | 8.161 | 85.37 | 0.000 |
| **Zebu cattle Only** |  |  |  |  |  |  |
| No Grouping | Among populations | 9 | 377.132 | 0.460 | 5.44 | 0.000 |
|  | Within populations | 730 | 5863.00 | 8.030 | 94.56 | 0.000 |
| Grouping-I (Provincial origin) (Group1-Tamil Nadu breeds; Group2-Andhra Pradesh breeds; Group3-Karnataka; Group4-Kerala) | Among groups | 3 | 134.12 | 0.027 | 0.32 | 0.326 |
|  | Among populations within groups | 6 | 243.00 | 0.443 | 5.21 | 0.000 |
|  | Within populations | 730 | 5863.00 | 8.032 | 94.47 | 0.000 |
| Grouping-II (Joshi and Phillips, 1953 classification) (Group1-ONG; Group2-DEO; Group3-ALA, BAR, HAL, KAN, UMB, PUL; GroupIV-VEC, PNR) | Among groups | 3 | 140.37 | 0.024 | 0.28 | 0.373 |
|  | Among populations within groups | 6 | 236.76 | 0.447 | 5.26 | 0.000 |
|  | Within populations | 730 | 5863.00 | 8.032 | 94.46 | 0.000 |
| Grouping III (Phylogeny) (Group 1-PUL, UMB; Group2-ALA, BAR; Group3-DEO, ONG, HAL; Group4-KAN; Group5-VEC; Group6-PNR) | Among groups | 5 | 228.17 | 0.101 | 1.19 | 0.088 |
|  | Among populations within groups | 4 | 148.97 | 0.375 | 4.41 | 0.000 |
|  | Within populations | 730 | 5863.00 | 8.032 | 94.41 | 0.000 |
| Grouping-IV (Genetic Structure) (Group1-DEO; Group2-ONG; Group3-BAR, ALA, HAL, PUL, UMB; Group4-VEC; Group5-PNR; Group6-KAN) | Among groups | 5 | 266.80 | 0.237 | 2.78 | 0.012 |
|  | Among populations within groups | 4 | 110.34 | 0.275 | 3.21 | 0.000 |
|  | Within populations | 730 | 5863.00 | 8.032 | 94.30 | 0.000 |

### 3.4. Mitochondrial DNA (mtDNA) control region diversity and haplogroup distribution

A 1207 bp long region of bovine mtDNA covering partial cytochrome B, t-RNA-Thr, t-RNA-Pro, D-loop (control region) and partial t-RNA-Phe coding DNA sequence were amplified in 523 South Indian cattle. After trimming and editing, the 951bp long final sequence corresponded to 15240 bp to 16190 bp of the complete mitochondrial genome (Hiendleder et al. 2008). All the sequences generated in the study were submitted to NCBI-GenBank and are available under accession numbers MW319823-MW320345. Overall, 144 polymorphic sites were observed in South Indian cattle sequences with 43 singletons and 101 parsimony informative sites. The overall nucleotide diversity and average number of nucleotide differences among South Indian cattle was 0.0111and 10.36 respectively. A total of 207 unique haplotypes were observed with overall haplotype diversity of 0.961 (Table 4). The haplotype diversity observed in most South Indian cattle breeds was relatively higher than that of North Indian zebu cattle located along the Gangetic plains (Sharma et al. 2015). Two major maternal haplogroups, I1 and I2, typical of zebu cattle were observed, with the former being predominant occurring 313 times and the later occurring 166 times. However, there were differences in certain breeds like Vechur and Umblachery in which I2 haplogroup was more frequent than I1. Interestingly, six zebu cattle showed taurine lineage, of which one belonged to T1 (Deoni), three belonged to T2 (one each from Hallikar, Bargur and Kangayam) and two belonged to T3 (one each from Deoni and Pulikulam). The overall nucleotide diversity within I1 and I2 haplogroups were 0.0024 and 0.0029 respectively while the overall average number of nucleotide differences were 2.32 and 2.77 respectively (Table 5). Among the 207 unique haplotypes, 105 belonged to I1 haplogroup, 74 belonged to I2 haplogroup, 3 belonged to T3 haplogroup and one each belonged to T1 and T2 halpogroups (Table 4). The overall haplotype diversity within I1 haplogroup and I2 haplogroup was 0.907 and 0.949 respectively. The haplotype diversity within I2 haplogroup was higher than that of I1 in all the South Indian zebu cattle breeds except Alambadi. Among the observed haplotypes, 27 were found to be shared across 2 or more breeds, of which 18 belonged to I1 haplogroup, 8 belonged to I2 haplogroup and one belonged to T3 haplogroup (Supplementary Table ST4). The haplotype HxJxABDLKOPUVHJ_H13 was the most frequent among I1 haplogroup and shared with all the investigated populations except Punganur cattle. Among the I2 haplogroup, HxJxABDLKOPUV_H7 was the most frequent haplotype and shared among 11 populations with the exception of Punganur, Holstein-Friesian and Jersey cattle. A total of 139 singleton haplotypes were observed, of which 67, 50, 18, 3 and one belonging to I1, I2, T3, T2 and T1 haplogroups respectively. The higher proportion of singleton haplotypes within I2 haplogroup of South Indian cattle was consistent with relatively higher haplotype and nucleotide diversity observed within this lineage.

**Table 4.** Haplogroup classification and diversity of mtDNA control region haplotypes in zebu, taurine and crossbred cattle of South India

| Breed | Haplogroup Count |  |  |  |  |  | Haplotype Count |  |  |  |  |  | Haplotype Diversity |  |  |  |
| --- | --- | --- | --- | --- | --- | --- | --- | --- | --- | --- | --- | --- | --- | --- | --- | --- |
|  | ALL | I1 | I2 | T1 | T2 | T3 | ALL | I1 | I2 | T1 | T2 | T3 | ALL | I1 | I2 | T3 |
| DEO | 53 | 39 | 12 | 1 | - | 1 | 30 | 18 | 10 | 1 | - | 1 | 0.952 | 0.914 | 0.970 | - |
| ONG | 51 | 29 | 22 | - | - | - | 20 | 9 | 11 | - | - | - | 0.914 | 0.778 | 0.913 | - |
| PNR | 16 | 14 | 2 | - | - | - | 7 | 5 | 2 | - | - | - | 0.692 | 0.593 | 1.000 | - |
| HAL | 36 | 22 | 13 | - | 1 | - | 28 | 16 | 11 | - | 1 | - | 0.981 | 0.957 | 0.974 | - |
| VEC | 26 | 10 | 16 | - | - | - | 10 | 5 | 5 | - | - | - | 0.892 | 0.756 | 0.800 | - |
| ALA | 29 | 21 | 8 | - | - | - | 20 | 14 | 6 | - | - | - | 0.970 | 0.952 | 0.929 | - |
| BAR | 52 | 34 | 17 | - | 1 | - | 24 | 16 | 7 | - | 1 | - | 0.939 | 0.882 | 0.890 | - |
| KAN | 51 | 36 | 14 | - | 1 | - | 18 | 11 | 6 | - | 1 | - | 0.882 | 0.784 | 0.835 | - |
| PUL | 31 | 19 | 11 | - | - | 1 | 21 | 11 | 9 | - | - | 1 | 0.972 | 0.942 | 0.956 | - |
| UMB | 33 | 16 | 17 | - | - | - | 19 | 10 | 9 | - | - | - | 0.955 | 0.900 | 0.912 | - |
| HFP | 15 | 10 | 1 | - | - | 4 | 11 | 6 | 1 | - | - | 4 | 0.905 | 0.778 | - | 1.000 |
| JER | 33 | 1 | - | - | - | 32 | 20 | 1 | - | - | - | 19 | 0.934 | - | - | 0.929 |
| HFX | 39 | 24 | 13 | - | - | 2 | 26 | 16 | 8 | - | - | 2 | 0.964 | 0.931 | 0.897 | 1.000 |
| JRX | 58 | 38 | 20 | - | - | - | 37 | 22 | 15 | - | - | - | 0.930 | 0.848 | 0.953 | - |
| Total/Overall | 523 | 313 | 166 | 1 | 3 | 40 | 207 | 105 | 74 | 1 | 3 | 24 | 0.961 | 0.907 | 0.949 | 0.940 |

**Table 5.** Polymorphism and diversity of mtDNA control region haplotypes in zebu, taurine and crossbred cattle of South India

| Breed | No. Polymorphic Sites |  |  |  | Singleton Sites |  |  |  | Parsimony Informative Sites |  |  |  | Nucleotide Diversity |  |  |  | Avg. No. Nucleotide Differences |  |  |  |
| --- | --- | --- | --- | --- | --- | --- | --- | --- | --- | --- | --- | --- | --- | --- | --- | --- | --- | --- | --- | --- |
|  | ALL | I1 | I2 | T3 | AL<br>L | I1 | I2 | T3 | ALL | I1 | I2 | T3 | ALL | I1 | I2 | T3 | ALL | I1 | I2 | T3 |
| DEO | 7 | 19 | 16 | - | 19 | 9 | 12 | - | 51 | 10 | 4 | - | 0.0072 | 0.0019 | 0.0036 | - | 6.86 | 1.83 | 3.39 | - |
| ONG | 22 | 22 | 15 | - | 4 | 4 | 7 | - | 18 | 18 | 8 | - | 0.0046 | 0.0014 | 0.0031 | - | 4.37 | 1.33 | 2.92 | - |
| PNR | 15 | 11 | 3 | - | 4 | 4 | 3 | - | 11 | 7 | 0 | - | 0.0038 | 0.0027 | 0.0032 | - | 3.58 | 2.52 | 3.00 | - |
| HAL | 67 | 19 | 14 | - | 47 | 13 | 9 | - | 20 | 6 | 5 | - | 0.0076 | 0.0027 | 0.0031 | - | 7.24 | 2.61 | 2.97 | - |
| VEC | 17 | 5 | 7 | - | 6 | 3 | 3 | - | 11 | 2 | 4 | - | 0.0047 | 0.0016 | 0.0021 | - | 4.51 | 1.51 | 1.98 | - |
| ALA | 26 | 14 | 9 | - | 13 | 8 | 7 | - | 13 | 6 | 2 | - | 0.0049 | 0.0026 | 0.0030 | - | 4.70 | 2.43 | 2.86 | - |
| BAR | 65 | 17 | 11 | - | 45 | 10 | 2 | - | 20 | 7 | 9 | - | 0.0064 | 0.0022 | 0.0029 | - | 6.10 | 2.10 | 2.79 | - |
| KAN | 58 | 11 | 9 | - | 40 | 4 | 5 | - | 17 | 7 | 4 | - | 0.0059 | 0.0018 | 0.0025 | - | 5.62 | 1.70 | 2.34 | - |
| PUL | 61 | 26 | 9 | - | 39 | 21 | 6 | - | 22 | 5 | 3 | - | 0.0086 | 0.0041 | 0.0025 | - | 8.18 | 3.87 | 2.38 | - |
| UMB | 24 | 11 | 10 | - | 8 | 5 | 5 | - | 16 | 6 | 5 | - | 0.0053 | 0.0025 | 0.0024 | - | 5.04 | 2.38 | 2.25 | - |
| HFP | 58 | 8 | - | 8 | 16 | 6 | - | 8 | 42 | 2 | - | 0 | 0.0211 | 0.0020 | - | 0.0042 | 20.00 | 1.91 | - | - |
| JER | 58 | - | - | 34 | 34 | - | - | 19 | 24 | - | - | 15 | 0.0066 | - | - | 0.4300 | 6.30 | - | - | 4.10 |
| HFX | 6 | 20 | 9 | 2 | 11 | 11 | 6 | 2 | 53 | 9 | 3 | 0 | 0.0094 | 0.0034 | 0.0021 | 0.0021 | 8.87 | 3.24 | 1.95 | 2.00 |
| JRX | 34 | 18 | 17 | - | 19 | 10 | 13 | - | 15 | 8 | 4 | - | 0.0045 | 0.0022 | 0.0027 | - | 4.26 | 2.04 | 2.58 | - |
| Overall | 144 | 83 | 67 | 41 | 43 | 38 | 31 | 25 | 101 | 45 | 36 | 16 | 0.0111 | 0.0024 | 0.0029 | 0.0042 | 10.36 | 2.32 | 2.77 | 3.97 |

### 3.5. mtDNA phylogeny and evolutionary relationship

Phylogenetic analysis was performed on all the 207 unique haplotypes along with reference sequences of each haplogroup, the results of which is presented in Supplementary Figure SF3. The phylogenetic tree conformed to haplogroup classification with the formation of two major indicine clusters I1 and I2. There were no major sub-clusters except for one minor clade containing four or more haplotypes in each of the I1 and I2 haplogroups. The minor clade within in I1 haplogroup was formed by four singleton haplotypes (K_H142, L_H120, L_H115 and V_H181) from Kangayam, Hallikar and Vechur cattle breed. Similarly, the minor clade within I2 haplogroup was formed by three singleton and two shared haplotypes (P_H155, AD_H61, B_H84, L_H129 and OP_H149) from Pulikulam, Alambadi, Deoni, Bargur, Hallikar and Ongole cattle.

The frequency of mtDNA control region haplotypes was further utilized to estimate pairwise genetic differentiation among zebu cattle from South India as well as the zebu inhabiting areas closer to Indus valley sites (West Indian zebu) and Gangetic plains (North Indian zebu and Nepalese zebu). Analysis of I1 haplotypes revealed pairwise F_ST_ varying between zero and o.293 while I2 haplotypes showed pairwise F_ST_ varying between zero and 0.255. The pairwise F_ST_ among I1 haplotypes was zero among NIC-NPC, WIC-NPC, ALA-UMB and HAL-UMB breed combinations and less than 1% among ALA-HAL, WIC-HAL, WIC-UMB, NIC-UMB, ALA-WIC breed combinations. The largest F_ST_ among I1 haplotypes was observed between Punganur cattle and other zebu breeds (Vechur, Ongole, Deoni, Bargur and Nepalese) followed by Vechur and other breeds (Pulikulam, Ongole and Umbalchery). In contrast, the pairwise F_ST_ among I2 haplotypes was zero between Punganur and other zebu breeds (ONG, BAR, NIC, UMB, DEO, HAL, WIC, ALA, KAN) and among HAL-DEO, WIC-DEO and PUL-DEO breed combinations. The largest F_ST_ among I2 haplotypes was observed between Nepalese zebu and South Indian zebu (ONG, KAN, PUL, BAR, UMB, VEC, DEO and ALA) followed by Vechur and other breeds (ALA, BAR, UMB, ONG, PUL, HAL and DEO).

The pairwise F_ST_ for each of the haplogroups were further utilized to perform principal components analysis (PCA) and understand the evolutionary relationships among the zebu cattle breeds (Figure 4). Principal components analysis (PCA) performs orthogonal transformation of a set of correlated variables (genetic distance estimations) into a set of new linearly uncorrelated variables called as principal components. The three largest principal components that explained maximum variation in the dataset were used to project sampled breeds into a three-dimensional geometric space. PCA of I1 haplotypes revealed the three largest principal components explaining 93.04% of the total variance in the dataset. The three-dimensional scattergram drawn using the three largest principal components showed the distinctness of Punganur, Pulikulam and Vechur cattle from all the other South Indian zebu cattle breeds. The zebu cattle from North India and Nepal were also found to be distinct from West and South Indian zebu cattle. Similarly, PCA of I2 haplotypes revealed the three largest principal components together contributing to 91.31% of total variance in the dataset. The three-dimensional scattergram showed Vechur and Nepalese cattle distinct from other zebu cattle breeds.

**Figure 4.**
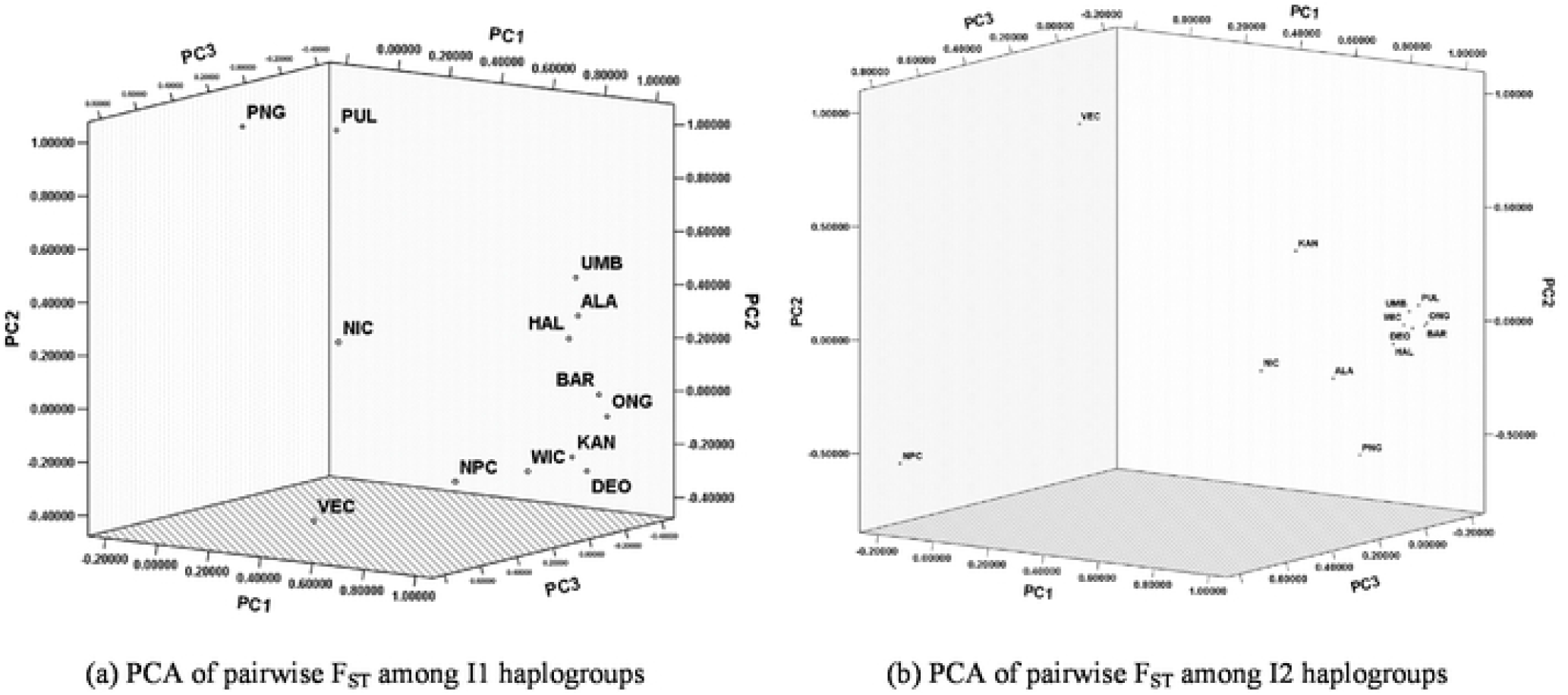
Scattergram of first three largest principal components derived from pairwise F_ST_ among mtDNA haplotypes of zebu cattle from South India, North India, West India and Nepal (ALA – Alambadi; BAR – Bargur; DEO – Deoni; HAL – Hallikar; KAN – Kangayam; ONG – Ongole; PNR – Punganur; PUL − Pulikulam; UMB – Umblachery; VEC – Vechur; NPC – Nepalese; NIC – North Indian; WIC – West Indian)

To understand the underlying genetic structure of each of the two maternal lineages (I1 and I2), analysis of molecular variance among zebu mtDNA haplotypes of cattle from South India, Gangetic plains (North India and Nepalese) and Indus valley (West India) sites were conducted (Table 6). When no grouping was assumed among I1 haplotypes, 6.17% of variance (P<0.05) was due to between population differences while 93.83% variance was due to within population differences. When the zebu cattle were grouped on the basis of their origin/location (South India, Gangetic Plains and Indus valley sites), among group variance was negligible (0.43%) and insignificant (P>0.05). However, when Punganur, Vechur and Pulikulam breeds were grouped separately from other South Indian zebu breeds, among group variance was 2.91% and statistically significant (P<0.05), thus confirming their genetic uniqueness. With respect to I2 haplotypes, when no grouping was assumed, among population variance was 8.32% (P<0.05) and within population variance was 91.61%. When grouped on the basis of their origin/location (South Indian, Gangetic Plains and Indus valley sites), among group variance was 2.24%, but not statistically significant (P>0.05). When Punganur, Vechur and Pulikulam breeds were grouped separately from other South Indian zebu breeds, among group variance reduced further to 2.01% (P>0.05). However, when Vechur and Nepalese cattle were grouped together, among group variance increased to 6.04% and was statistically significant (P<0.05). Thus, the mtDNA control region variation of both I1 and I2 lineages showed Vechur cattle as distinct from other South Indian breeds while Punganur and Pulikulam cattle were also distinct among the I1 maternal lineage. Vechur and Punganur are dwarf cattle breeds and morphologically distinct from rest of the South Indian zebu while Pulikulam is primarily a draught breed, compact in size and mainly used for draught and penning in agricultural fields.

**Table 6.**
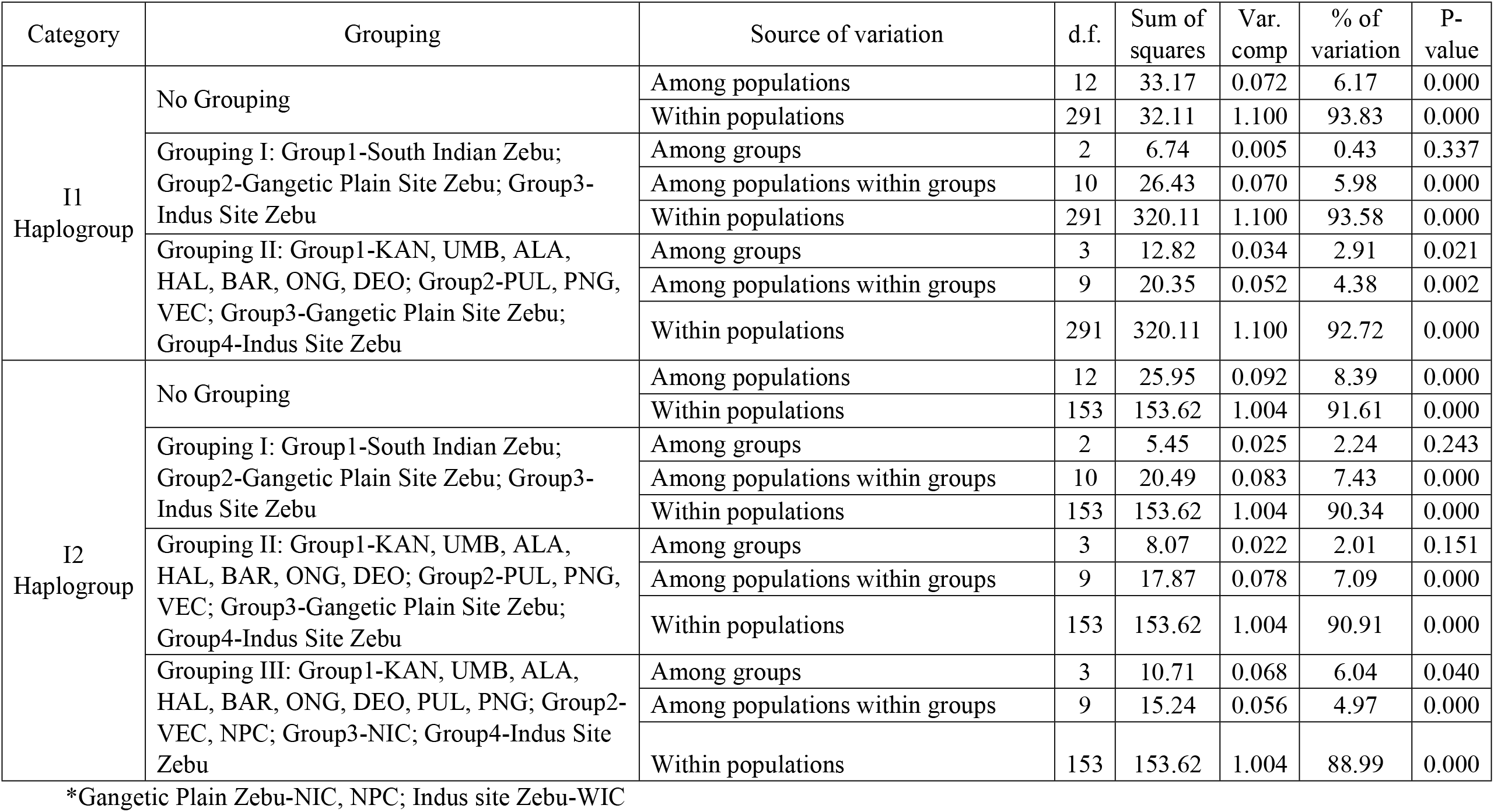
Analysis of molecular variance among mtDNA haplotypes of zebu cattle of South India, Indus valley site and Gangetic plains

| Category | Grouping | Source of variation | d.f. | Sum of squares | Var. comp | % of variation | P-value |
| --- | --- | --- | --- | --- | --- | --- | --- |
| I1<br>Haplogroup | No Grouping | Among populations | 12 | 33.17 | 0.072 | 6.17 | 0.000 |
|  |  | Within populations | 291 | 32.11 | 1.100 | 93.83 | 0.000 |
|  | Grouping I: Group1-South Indian Zebu;<br>Group2-Gangetic Plain Site Zebu; Group3-Indus Site Zebu | Among groups | 2 | 6.74 | 0.005 | 0.43 | 0.337 |
|  |  | Among populations within groups | 10 | 26.43 | 0.070 | 5.98 | 0.000 |
|  |  | Within populations | 291 | 320.11 | 1.100 | 93.58 | 0.000 |
|  | Grouping II: Group1-KAN, UMB, ALA, HAL, BAR, ONG, DEO; Group2-PUL, PNG, VEC; Group3-Gangetic Plain Site Zebu; Group4-Indus Site Zebu | Among groups | 3 | 12.82 | 0.034 | 2.91 | 0.021 |
|  |  | Among populations within groups | 9 | 20.35 | 0.052 | 4.38 | 0.002 |
|  |  | Within populations | 291 | 320.11 | 1.100 | 92.72 | 0.000 |
| I2<br>Haplogroup | No Grouping | Among populations | 12 | 25.95 | 0.092 | 8.39 | 0.000 |
|  |  | Within populations | 153 | 153.62 | 1.004 | 91.61 | 0.000 |
|  | Grouping I: Group1-South Indian Zebu;<br>Group2-Gangetic Plain Site Zebu; Group3-Indus Site Zebu | Among groups | 2 | 5.45 | 0.025 | 2.24 | 0.243 |
|  |  | Among populations within groups | 10 | 20.49 | 0.083 | 7.43 | 0.000 |
|  |  | Within populations | 153 | 153.62 | 1.004 | 90.34 | 0.000 |
|  | Grouping II: Group1-KAN, UMB, ALA, HAL, BAR, ONG, DEO; Group2-PUL, PNG, VEC; Group3-Gangetic Plain Site Zebu; Group4-Indus Site Zebu | Among groups | 3 | 8.07 | 0.022 | 2.01 | 0.151 |
|  |  | Among populations within groups | 9 | 17.87 | 0.078 | 7.09 | 0.000 |
|  |  | Within populations | 153 | 153.62 | 1.004 | 90.91 | 0.000 |
|  | Grouping III: Group1-KAN, UMB, ALA, HAL, BAR, ONG, DEO, PUL, PNG; Group2-VEC, NPC; Group3-NIC; Group4-Indus Site Zebu | Among groups | 3 | 10.71 | 0.068 | 6.04 | 0.040 |
|  |  | Among populations within groups | 9 | 15.24 | 0.056 | 4.97 | 0.000 |
|  |  | Within populations | 153 | 153.62 | 1.004 | 88.99 | 0.000 |

\*Gangetic Plain Zebu-NIC, NPC; Indus site Zebu-WIC

### 3.6. Mismatch distribution, haplotype network and tests for selective neutrality

To obtain further insight into the demographic history of South Indian draught type zebu cattle, the mismatch distribution of pairwise nucleotide differences among haplotypes within I1 and I2 maternal lineages was analysed for each of the investigated breeds (Figure 5). Sudden or exponential demographic population expansion/decline are characterized by unimodal distribution and the mismatch values increase over time as mutations accumulate in a population. In contrast, stable populations show a “ragged” multimodal distribution as deep divergences accumulate between lineages (Rogers and Harpending 1992, Schenekar and Weiss, 2011; Grant 2015). With respect to I1 lineage, unimodal distribution was observed in most of the South Indian breeds with the modal value at two pairwise mismatches. Bimodal distribution was observed among I1 haplotypes in Kangayam, Umbalchery and Punganur breeds with modal values at 1 and 3, 1and 5, 2 and 6 mismatches respectively. In case of zebu cattle from Indus valley site (WIC), the modal value was observed at 3 mismatches while the Gangetic plains Zebu cattle, NIC and NPC showed modal values at 3 and 2 mismatches respectively. The modal values of South Indian I1 lineage observed in the present study is contrary to the reports made earlier (Chen et al. 2010). Similarly, with respect to I2 lineage, modal values were observed around two to four mismatches in most of South Indian and Gangetic Plain Zebu cattle (NIC and NPC) while it was observed at one mismatch in Indus valley site zebu cattle (WIC). Unimodal distribution of I2 haplotypes was observed in most South Indian zebu breeds except Deoni, Kangayam and Pulikulam. The mismatch analysis of pairwise differences revealed the parameter ‘tau’ (the coalescent time of expansion) ranging from 1.268 (Ongole) to 8.406 (Punganur) among I1 haplotypes while it ranged from 1.344 (Pulikulam) to 4.182 (Kangayam) among I2 haplotypes of South Indian zebu cattle. The estimated sum of squared deviations (SSD) and the raggedness index was significant (P<0.05) in Deoni and Bargur cattle for I1 and I2 haplotypes respectively (Table 7).

**Table 7.** Tests for demographic expansion based on mismatch analysis and neutrality in South Indian draught type Zebu cattle

| Statistics | ALA | BAR | DEO | HAL | KAN | ONG | PNG | PUL | UMB | VEC |
| --- | --- | --- | --- | --- | --- | --- | --- | --- | --- | --- |
| **Mismatch analysis (I1 haplogroup)** |  |  |  |  |  |  |  |  |  |  |
| Tau | 2.221 | 1.693 | 1.709 | 1.906 | 1.818 | 1.268 | 8.406 | 2.365 | 1.963 | 2.186 |
| SSD | 0.015 | 0.001 | 0.020 | 0.005 | 0.004 | 0.008 | 0.077 | 0.014 | 0.009 | 0.034 |
| Model (SSD) p-value | 0.090 | 0.850 | 0.030 | 0.630 | 0.650 | 0.280 | 0.270 | 0.430 | 0.410 | 0.240 |
| Raggedness index | 0.091 | 0.029 | 0.134 | 0.070 | 0.041 | 0.106 | 0.230 | 0.041 | 0.084 | 0.135 |
| Raggedness p-value | 0.130 | 0.910 | 0.020 | 0.350 | 0.790 | 0.160 | 0.350 | 0.820 | 0.270 | 0.430 |
| **Mismatch analysis (I2 haplogroup)** |  |  |  |  |  |  |  |  |  |  |
| Tau | 2.111 | 2.055 | 1.969 | 1.832 | 4.182 | 1.891 | - | 1.344 | 1.777 | 2.438 |
| SSD | 0.008 | 0.032 | 0.018 | 0.003 | 0.135 | 0.003 | - | 0.042 | 0.007 | 0.014 |
| Model (SSD) p-value | 0.900 | 0.030 | 0.300 | 0.870 | 0.170 | 0.830 | - | 0.110 | 0.480 | 0.460 |
| Raggedness index | 0.082 | 0.163 | 0.129 | 0.070 | 0.403 | 0.029 | - | 0.269 | 0.078 | 0.049 |
| Raggedness p-value | 0.770 | 0.020 | 0.180 | 0.590 | 0.150 | 0.970 | - | 0.070 | 0.360 | 0.820 |
| **Neutrality Tests (I1 haplogroup)** |  |  |  |  |  |  |  |  |  |  |
| Tajima's D | -1.015 | -1.283 | -1.780 | -1.718 | -1.031 | -1.102 | -0.813 | -1.565 | -0.559 | -0.582 |
| Tajima's D p-value | 0.139 | 0.074 | 0.017 | 0.029 | 0.150 | 0.133 | 0.214 | 0.040 | 0.302 | 0.308 |
| Fu's FS | -27.168 | 27.892 | -27.627 | -27.106 | -27.594 | -29.071 | -17.187 | -26.340 | -23.739 | -12.311 |
| FS p-value | 0.000 | 0.000 | 0.000 | 0.000 | 0.000 | 0.000 | 0.000 | 0.000 | 0.000 | 0.000 |
| **Neutrality tests (I2 haplogroup)** |  |  |  |  |  |  |  |  |  |  |
| Tajima's D | -1.170 | -0.633 | -1.705 | -1.770 | -0.908 | -1.307 | 0.000 | -1.465 | -1.141 | 0.258 |
| Tajima's D p-value | 0.142 | 0.309 | 0.035 | 0.022 | 0.220 | 0.088 | 100.000 | 0.048 | 0.127 | 0.648 |
| Fu's FS | -7.179 | -26.354 | -14.470 | -17.112 | -19.983 | -27.790 | 0.000 | -17.081 | -26.857 | -24.732 |
| FS p-value | 0.000 | 0.000 | 0.000 | 0.000 | 0.000 | 0.000 | 0.249 | 0.000 | 0.000 | 0.000 |

**Figure 5.**
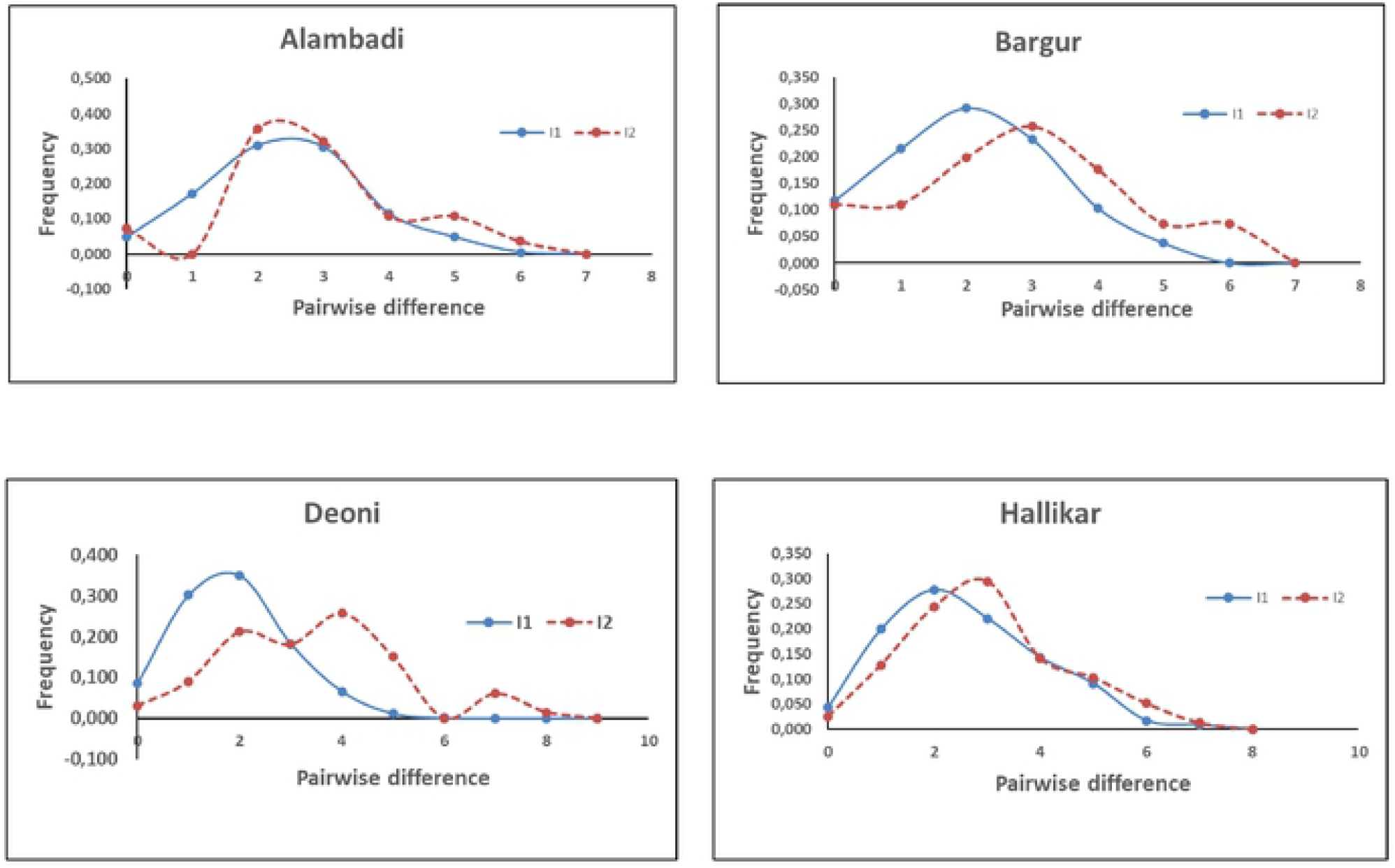
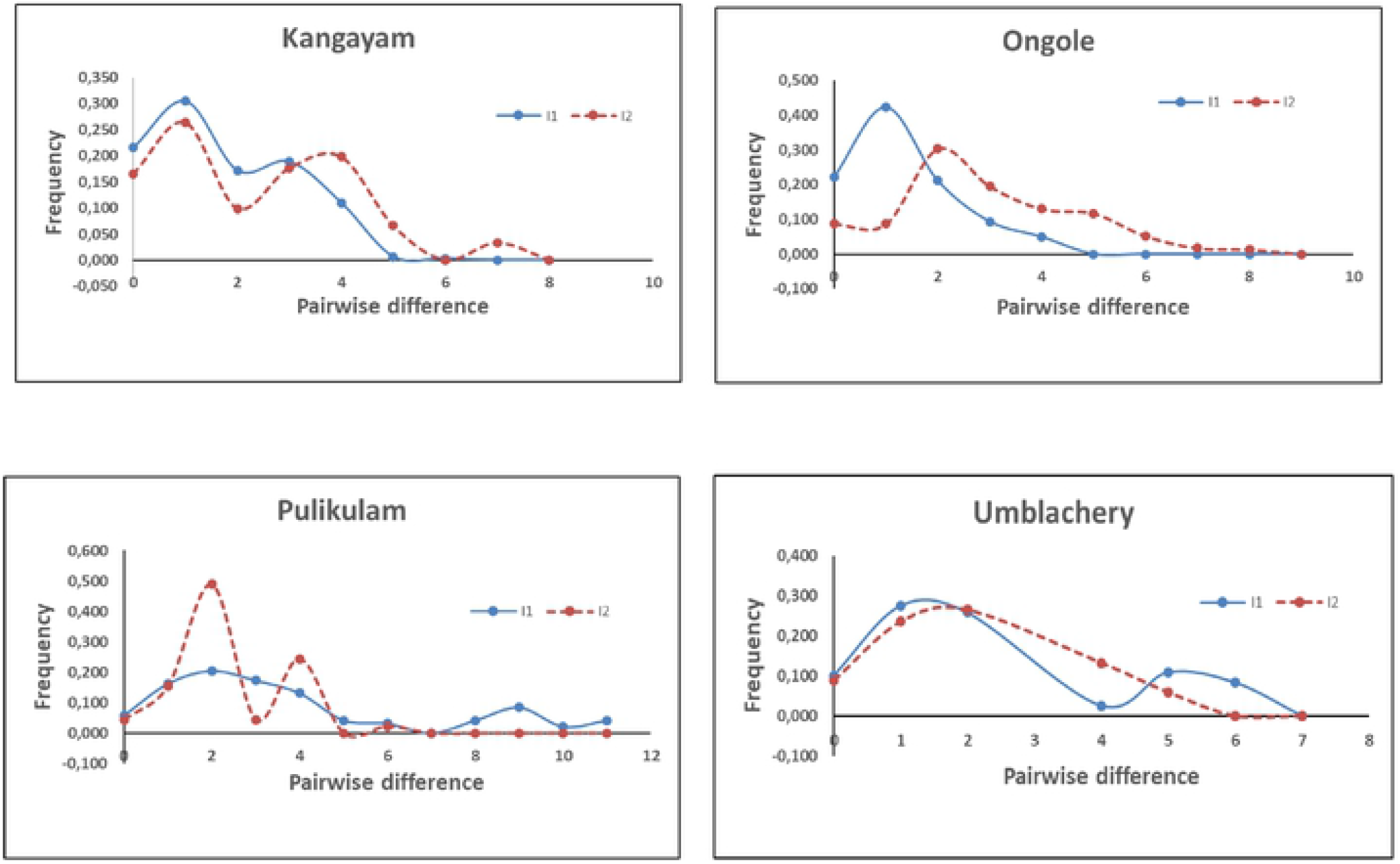
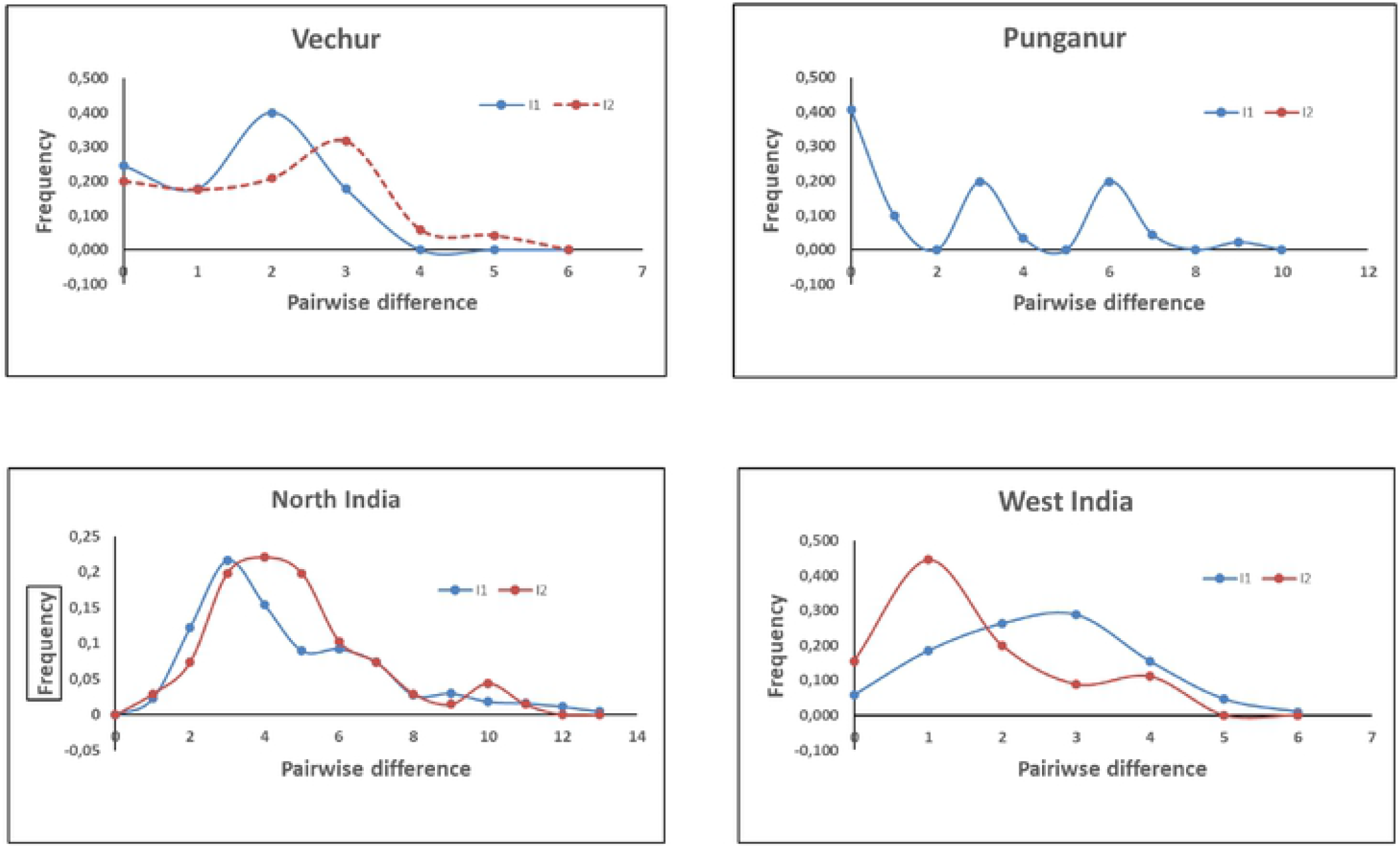
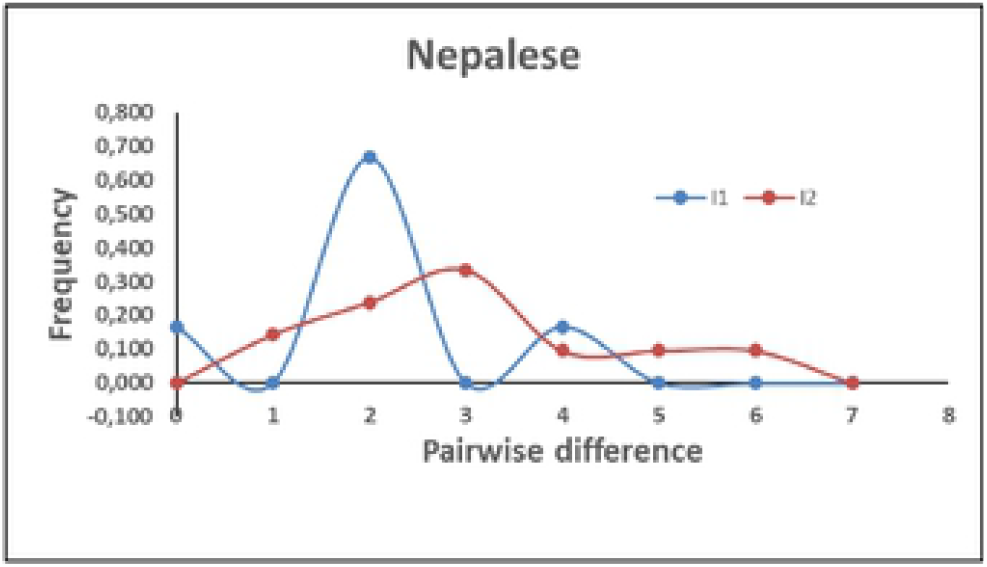
Mismatch distribution of pairwise nucleotide differences among mtDNA control region haplogroups of zebu cattle

Median Joining (MJ) network of mtDNA haplotypes from South Indian zebu cattle was reconstructed. Additionally, 298 mtDNA control region sequences of North Indian and Nepalese (Gangetic Plains), West Indian (Indus valley site), Chinese, Bhutanese and Vietnamese cattle were also included for haplotype reconstruction. MJ network reconstruction focused on loci with substitution mutations and did not consider loci with indels for the analysis. The network results (Figure 6) revealed two distinct lineages, I1 and I2 as expected with two star like expansion events radiating from ancestral nodes. Each of these two ancestral nodes were fed by singleton or low frequent haplotypes observed in zebu cattle breeds from Southern India, Gangetic plains and Indus valley sites. An additional star like expansion event of smaller magnitude was observed within the I1 haplogroup indicating a possible sub-lineage of I1. The South Indian zebu cattle were further subjected to tests for selective neutrality, the Tajima’s D and Fu’s FS. The Tajima’s D statistics was based on frequency spectrum of mutations while Fu’s FS was based on haplotype distribution. The results revealed negative Tajima’s D in all the investigated South Indian breeds for haplogroup I1, of which Deoni, Hallikar and Pulikulam were statistically significant (P<0.05). Similarly, Fu’s FS was negative and highly significant (P<0.01) for all the studied breeds. In case of I2 haplogroup, Tajima’s D was negative in all the studied breeds except Vechur cattle, of which Deoni, Hallikar and Pulikulam breeds were statistically significant (P<0.05). The Fu’s FS was negative and highly significant (P<0.01) for I2 haplogroups in all South Indian cattle breeds except Punganur. The negative Tajima’s D signifies an excess of low frequency polymorphisms than expected, indicating either expansion of population size or purifying selection. Similarly, negative FS is evidence for an excess number of alleles as would be expected from a recent population expansion or genetic hitchhiking. The results of the present study thus indicated purifying selection among the South Indian zebu cattle breeds. Further, Fu’s FS statistic, sensitive to the presence of singletons and devised more specifically to detect demographic expansion was statistically significant (P<0.01) in all the investigated cattle breeds.

**Figure 6.**
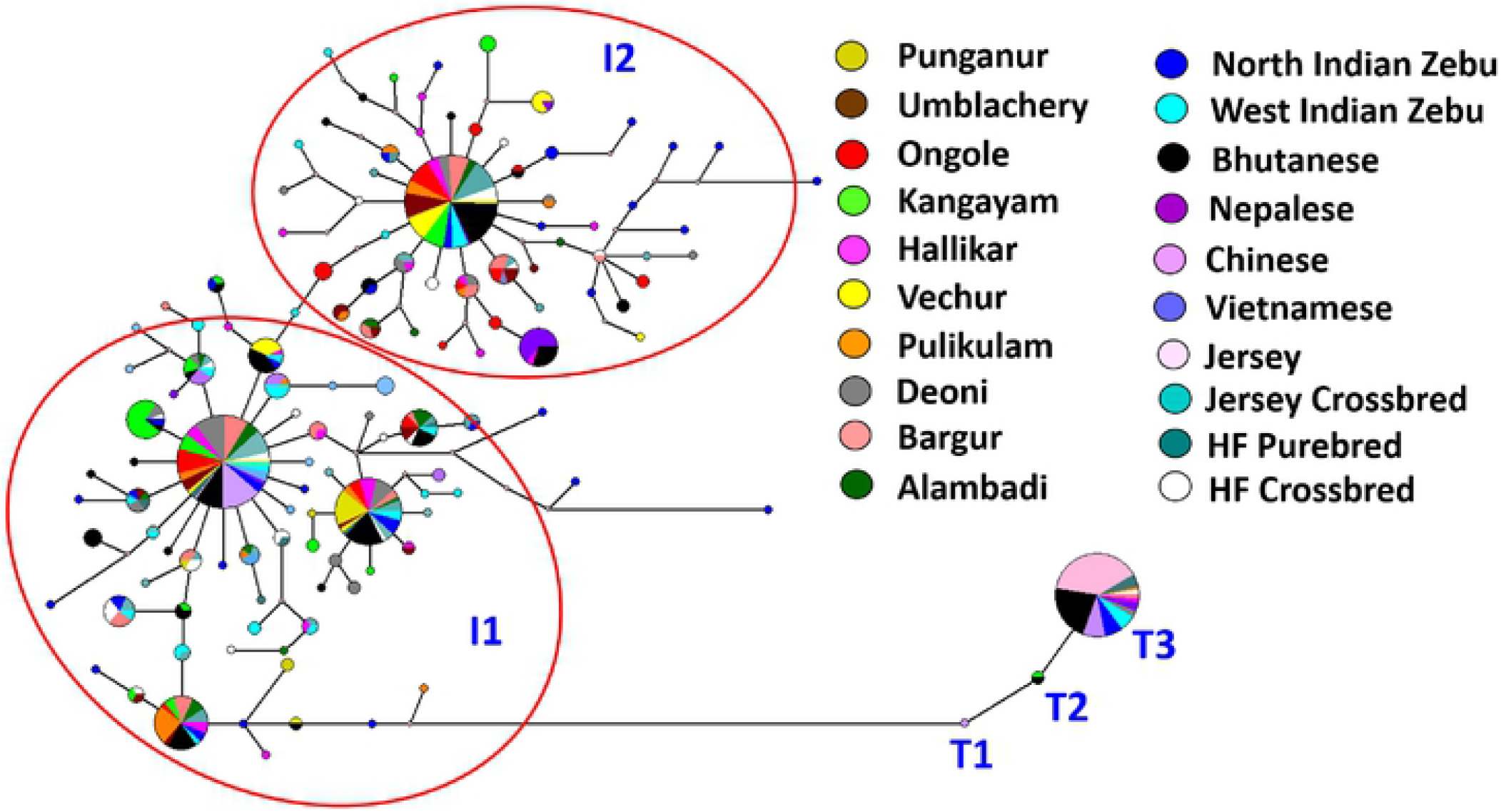
Median Joining network of mtDNA control region haplotypes of draught type zebu, taurine and crossbred cattle from South India and Zebu cattle from China, Nepal, Vietnam, North India, West India (Size of the circle indicates frequency of haplotypes)

## 4. DISCUSSION

Our study presents the first comprehensive genetic and phylogeographic analysis of draught type zebu cattle breeds of South India using nuclear and mitochondrial DNA variations. Historically, the focus of cattle breeding in South India was towards improving draught characteristics, mainly for agricultural operations and transport, including movement of military equipment. However, a wide diversity of zebu cattle breeds exists in South India with body size ranging from dwarf (Vechur, Punganur) to compact (Hallikar, Bargur, Umblachery, Alambadi, Ongole, Pulikulam) and medium to large (Deoni, Kangayam) size animals; draught characteristics varying from heavy load pulling capability (Hallikar, Umblachery) to speed and endurance in trotting (Kangayam, Bargur); utilities ranging from draught to penning for dung (Pulikulam, Bargur). Joshi and Phillips (1953) classified Zebu cattle of India and Pakistan into six major groups based on their morphology, breed history and phenotypic characteristics. The South Indian zebu cattle investigated in the present study were classified under three major categories: Group II, Short horned, white or light grey cattle with coffin shaped skulls (Ongole); Group III, Ponderously, built Red and White cattle (Deoni); and Group IV, Mysore type, compact sized cattle (Hallikar, Kangayam, Umblachery, Bargur, Pulilkulam and Alambadi). The bantam weight dwarf type cattle breeds, Vechur and Punganur did not fall into any of these three categories.

### 4.1. Population status, effective population size and inbreeding estimates

With rapid mechanization of agriculture in South India, the need for rearing draught cattle came down drastically in the last few decades. This led to reduction in effective population size and consequent increase in rate of inbreeding in many of these breeds (Singh and Sharma, 2017). A significantly strong negative Pearson’s correlation coefficient of −0.674 (P<0.05) was observed between the effective population size of different breeds and their estimated F_IS_ in the present study. The relatively higher mean F_IS_ estimates of some of the breeds reflected the breedable male and female populations (DADF, 2015) in their respective native breed tracts. For example, huge difference in the population of breedable male vs breedable female (25 vs 2977) of Pulikulam cattle resulted in low effective population size (adjusted Ne = 69). The highest estimate of F_IS_ among the breeds investigated in the present study was observed in Pulikulam cattle (F_IS_=0.108). Vechur cattle with second highest estimated F_IS_ (0.089) had only few hundred breedable females and <5 breedable males with an effective population size of <10. Similarly, Umblachery and Bargur breeds whose effective population size was <1600, had higher mean F_IS_ estimates of 0.089 and 0.076 respectively. Although the total population of both these breeds were relatively higher (39050 and 14154 heads of Umblachery and Bargur respectively), only few hundred breedable males (586 and 624 for Umblachery and Bargur respectively) were present in their respective native breed tracts. Lack of availability of sufficient breeding bulls was one of the main reasons for low effective population size, higher estimates of F_IS_ and high rate of inbreeding in these breeds (Singh and Sharma, 2017). Among the South Indian zebu cattle, the lowest F_IS_ estimate was observed in Hallikar cattle (F_IS_=0.021), whose total population was greater than 1.2 million heads with breedable male and female population of 5524 and 397357 respectively. Interestingly, Punganur cattle with a total population of less than 3000 and effective population size of ∼200 showed positive, but relatively lower estimate of F_IS_ (0.041). This could be possibly due to a better breedable male to breedable female ratio observed in this breed (0.0715) as compared to Vechur (0.0020), Pulikulam (0.0084) and Umbalchery (0.0387) cattle (DADF, 2015).

Singh and Sharma (2017) assessed the degree of endangerment of Indian zebu cattle breeds based on estimated breed-wise cattle population data (DADF, 2015), effective population size and estimated rate of inbreeding. Using the criteria laid down by Food and Agriculture Organization of the United Nations (FAO, 2013) and the Indian Council of Agricultural Research-National Bureau of Animal Genetic Resources (ICAR-NBAGR, 2016), they classified the status of Vechur as “critical” (breeding males less than five, breeding females 494 and high rate of inbreeding), Punganur and Pulikulam as “endangered” (breeding females more than 300 but less than 3000) and Bargur as “vulnerable” (breeding females more than 3000 but less than 6000). Thus, the status of four out of ten South Indian zebu cattle breeds investigated in the present study are in the category of “under risk” from conservation standpoint. The level of inbreeding estimates is consistent with their population status and is a cause for concern from conservation standpoint. It is also worth to mention that the population status of “Alambadi” breed is currently not known but has been observed to retain moderate levels of heterozygosity.

### 4.2. Origin, breed history and population structure

Most South Indian cattle breeds belong to Mysore type cattle (Group IV), of which Hallikar is considered to be the ancient one and widely distributed with significant and stable population size. Ware (1942) and Joshi and Phillips (1953) mentioned the origin of most Mysore type cattle breeds centred around Hallikar cattle. The results of AMOVA and genetic structure analysis obtained in the present study was in conformity with this theory of origin for most, if not all the breeds. Among group variance was significant (P<0.05) when all the Mysore type breeds were grouped together with the exception of Kangayam. Estimation of fixation index showed the pairwise F_ST_ was lowest among Hallikar and Alambadi (F_ST_=0.023), Alambadi and Bargur (F_ST_=0.026), Alambadi and Pulikulam (F_ST_=0.024), Bargur and Umblachery (F_ST_=0.031) breeds. Further, significant level of Hallikar ancestry was observed in Alambadi, Bargur, Pulikulam, and Umblachery cattle breeds (Figure 3).

Alambadi cattle are bred in the hilly tracts of Western ghats (Salem and Erode districts of the state of Tamil Nadu) and are raised mostly by grazing in the forest areas (Vandana et al. 2020). These cattle, known for their draught characteristics have close resemblance to Hallikar that are reared in the adjoining province of erstwhile Mysore. Ware (1942) expressed doubt whether Alambadi cattle need to be considered as a pure breed or an offshoot of the Hallikar. Although Bayesian clustering analysis showed Hallikar ancestry in Alambadi cattle, subsequent admixture of Bargur cattle was observed. This is understandable, considering the geographical overlap of the native tracts of both the breeds (Alambadi and Bargur) and the possibility of gene flow while grazing deep inside the forest areas of Western ghats. Bargur, the hill cattle of Tamil Nadu, are smaller and more compact, with not so prominent forehead as observed in other Mysore type breeds. They have distinct morphological appearance due to their coat colour, typical of being red and white or red with white spots. These cattle are known for their speed and endurance in trotting, but are very fiery, restive and difficult to train (Ware, 1942; Ganapathi et al. 2009; Pundir et al 2009). It also needs to be noted that among these four Mysore type Tamil Nadu breeds (Alambadi, Bargur Umblachery and Pulikulam), significant selection process had gone into the development of Bargur cattle, particularly for their unique morphological characteristics, herd uniformity and adaptation to zero input, hilly environment. This was evidenced by their individual assignment to unique inferred cluster when K=7 to K=8 (Figure 3) was assumed under Bayesian clustering analysis.

Pulikulam, is a migratory cattle breed, maintained in a zero-input system in southern parts of Tamil Nadu state of South India. These cattle have predominantly white or greyish coat colour and are mostly used for agricultural operations and rural transport. These cattle are well-known for their utilities as a sport animal for Jallikattu (a traditional bull embracing sport) and penning (a practice of manuring by keeping the cattle overnight in open agricultural fields). These cattle resemble Kangayam in physical appearance but are significantly smaller and compact in size. Gunn (1909) opined that these cattle probably had a strain of Mysore cattle blood in them. The present study revealed Pulikulam as a genetically distinct breed from Kangayam but sharing significant ancestry with Hallikar and Deoni cattle (Figure 3; K=4 to K=7). The genetic difference between Pulikulam and Kangayam was also consistent with the reported differences in morphometric parameters that included body length, height at withers and chest girth (Kanakraj and Kathiresan, 2006; Singh et al. 2012).

Umblachery is an excellent draught cattle breed reared in the eastern coastal areas of Tamil Nadu state (Rajendran et al. 2008). These are medium sized cattle, selectively bred for short stature and suitability to work in marshy paddy fields. Anecdotal evidence attributed the possible origin and development of Umblachery from Kangayam cattle. Gunn (1909) mentioned that with the exception of appearance of head of these animals due to dehorning, these cattle retained the chief characteristics of Kangayam. However, the present study showed significant genetic differentiation among these breeds (F_ST_=0.070) and little or no traces of Kangayam ancestry in Umblachery cattle. Similar to Pulikulam cattle, significant admixture was observed in Umblachery cattle with traces of Hallikar and Deoni ancestry in them. This could be possibly due to trade influx of cattle into native tract of Umbalchery more than a century ago, from Southern Tamil Nadu (Cumbum district) where the now extinct breed called “Kappiliyan” or “Tambiran Madu” of Canarese (Mysore) origin was in abundance (Gunn, 1909; Littlewood, 1936).

The genetic structure analysis clearly indicated the influence of Hallikar breed in the development of most Mysore type cattle breeds of Tamil Nadu (Alambadi, Bargur, Umblachery and Pulikulam) with the exception of Kangayam. Kangayam is one of the popular draught breeds of South India known for their power (0.8hp per pair of bullocks) and sturdiness (Surendrakumar, 1988). Although, there are reports indicating the existence of Kangayam cattle for nearly three centuries (Reddi, 1957), the origin and development of this breed has been attributed mainly to 33^rd^ Pattagar (community chief) of Palayakottai who selectively bred the local cattle, only with sires available in his herd. The appearance of some of his cows/heifers gave the impression that they had a distinct strain of Ongole blood in them, as the Pattagar was known to possess purebred Ongole cows (Gunn, 1909). But no records were maintained by Pattagar during the early stages of the development of this breed to confirm this. Kangayam cattle conform largely to Mysore type breeds, but with a relatively larger body size, probably indicating the admixture with a heavier breed (Phillips, 1944). However, the results of phylogeny, multidimensional scaling and analysis of molecular variance in the present study clearly showed the genetic uniqueness of Kangayam cattle. The pairwise F_ST_ indicated marked differentiation of Kangayam from Hallikar (F_ST_=0.093) and Ongole (F_ST_=0.075) cattle. Further, Kangayam was also significantly differentiated from other Mysore type breeds (Alambadi, Bargur, Umblachery and Pulikulam) with pairwise F_ST_ ranging from 6.3% to 7.7%. The Bayesian clustering analysis showed Kangayam as a genetically distinct entity from both Ongole and Hallikar cattle. Low levels of admixture observed in Kangayam cattle, when assumed under K=4 to K=8 (Figure 3), was probably due to ancestral zebu inheritance common to all South Indian breeds. The genetic uniqueness of Kangayam cattle might have been the outcome of systematic selective breeding practices of Pattagars initially, and subsequent introduction of herd book registration followed by implementation of various genetic improvement programs since1940s by Government of Madras (Panneerselvam and Kandasamy, 2008).

The two bantam cattle breeds, Vechur and Punganur are unique to South India and are probably among the smallest cattle breeds of the world, with the average height of adults ranging from 60-100 cm and body weight of 125-200 kg (Nath, 1993; Iype, 1996). Vechur and Punganur cattle have retained moderate levels of diversity despite the small population size and being classified in the critically endangered category. Both these breeds were distinct from rest of the South Indian breeds, not only in terms of morphology but also in terms of milk production efficiency. The cows are considered to be efficient milk producers relative to their size, with peak milk yield scaling up to 3-4 kg/day (Nath, 1993; Iype, 1996). Genetic distance based phylogenetic analysis and multidimensional scaling plot derived from pairwise F_ST_ showed Vechur and Punganur to be genetically distinct from other South Indian draught type zebu cattle. Although, Bayesian clustering analysis indicated the influence of Hallikar ancestry in both the breeds (K=3 to K=6), they clustered independently at K=7 and K=8 (Figure 3). It is also worth to mention that a high degree of genetic differentiation was observed among these two dwarf cattle (Vechur and Punganur) breeds (F_ST_=0.092). This is understandable considering the geographical location of their respective native tracts. The isolation of native tract of Vechur cattle (Kottayam district of Kerala state) by rivers, canals and backwaters and the distant location of Punganur cattle habitat (Chittoor district of Andhra Pradesh state) coupled with selective breeding practiced by farmers, might have contributed to the genetic differentiation of the two breeds. In addition to restricted gene flow, genetic drift arising out of small population size of both the breeds could have also contributed to their genetic divergence.

### 4.3. Breed purity and level of taurine introgression

With the mechanization of agricultural operations and increasing demand for dairy products in the last few decades, the focus of cattle breeding in South India shifted towards improving milk production. Beginning with the Operation Flood program in 1970s, intensive cattle development projects in Southern India were targeted towards genetic improvement of local cattle for milk through crossbreeding with exotic commercial taurine cattle. Although the official breeding policy emphasized the need for selective breeding of recognized zebu cattle breeds, indiscriminate crossbreeding took place in the native tracts of many indigenous breeds. In the absence of pedigree recording under small holder production set up, it is not uncommon that the local cows inseminated with purebred taurine or crossbred semen were subsequently served by locally available sires or *vice versa*. Often, the choice of semen used for artificial insemination (AI) is left to the inseminators and depends on the availability of purebred zebu, purebred taurine or crossbred semen straws, irrespective of the breed/genotype of the cows (purebred zebu, non-descript local, crossbred cattle, etc.) brought for AI. This resulted in dilution of purebred zebu germplasm with varying levels of taurine admixture in well described zebu cattle populations, thus affecting their breed purity.

Bayesian clustering analysis was performed to assess the taurine admixture in South Indian zebu cattle using purebred Jersey and Holstein-Friesian as reference genotypes. Samples from *Bos taurus X Bos indicus* crossbreds were also genotyped to evaluate the range of taurine inheritance under field conditions. The proportion of membership coefficient obtained for individual animals in the inferred zebu and taurine clusters were utilized to estimate taurine admixture and purity of the investigated zebu cattle breeds (Table 8). Ongole had zero individuals with >6.25% taurine admixture while Punganur had the highest proportion (44.4%) of individuals with >6.25% taurine admixture. Among the South Indian cattle, Ongole, Kangayam, Deoni, Alambadi and Bargur breeds showed majority of their individuals (>95% of each of these populations) with less than 6.25% taurine admixture. The relatively low level of taurine admixture in Ongole, Kangayam and Deoni can be attributed to the following factors: (i) better artificial insemination coverage in their respective breed tracts (ii) availability of purebred zebu, true to the type breeding bulls (iii) availability of purebred zebu semen for AI and (iv) better implementation of breeding policy in favour of selective breeding of purebred zebu cattle (Panneerselvam and Kandasamy, 2008; Natarajan et al. 2012; Dongre et al. 2017). In case of Alambadi and Bargur cattle raised in a unique, extensive production system of hilly areas, the access to artificial insemination services is little or absent. The cows in heat are normally served by the bulls that accompany the herd while grazing in the forest areas. For example, in Bargur cattle, the herd composition includes at least 2% of total size as breeding bulls and young male stocks (Ganapathi et al. 2009) in addition to 18-20% bullocks. The farmers select the breeding bulls based on certain phenotypic characteristics including uniformity of coat colour (Pundir et al 2009). The absence of AI facility and natural service being the main breeding practice in the native tract, the introduction of exotic bull semen was limited in Bargur and Alambadi cattle resulting in low levels of taurine admixture in these breeds. With respect to Hallikar cattle, the proportion of individuals with >6.25% taurine admixture was estimated to be 16.7%. This was surprisingly higher, considering the significant and stable population size of this breed (DADF, 2015). Singh et al. (2008) reported the demand for purebred Hallikar semen was higher than the available supply in the native tract and thus the lack of quality breeding bulls might have contributed to increased levels of taurine admixture in this breed.

**Table 8.** Overall breed wise proportion of membership coefficient in inferred taurine and zebu clusters and proportion of cattle with different levels of taurine admixture

| Lab | N | Overall proportion of membership coefficient in inferred clusters |  | Proportion of cattle (in %) with different levels of taurine admixture |  |  |  |
| --- | --- | --- | --- | --- | --- | --- | --- |
|  |  | Taurine | Zebu | >6.25% | >12.5% | >25% | >50% |
| ALA | 27 | 0.014 | 0.986 | 3.7 | 0 | 0 | 0 |
| BAR | 50 | 0.016 | 0.984 | 4 | 0 | 0 | 0 |
| HAL | 36 | 0.042 | 0.958 | 16.7 | 13.9 | 2.8 | 0 |
| KAN | 50 | 0.011 | 0.989 | 2 | 0 | 0 | 0 |
| PUL | 34 | 0.059 | 0.941 | 23.5 | 14.7 | 5.9 | 0 |
| UMB | 33 | 0.074 | 0.926 | 33.3 | 21.2 | 6.1 | 3.0 |
| DEO | 47 | 0.015 | 0.985 | 2.1 | 0 | 0 | 0 |
| ONG | 49 | 0.008 | 0.992 | 0 | 0 | 0 | 0 |
| VEC | 26 | 0.072 | 0.928 | 42.3 | 26.9 | 3.8 | 0 |
| PNR | 18 | 0.101 | 0.899 | 44.4 | 22.2 | 11.1 | 5.6 |
| HFP | 15 | 0.948 | 0.052 | 100 | 100 | 100 | 100 |
| JER | 34 | 0.991 | 0.009 | 100 | 100 | 100 | 100 |
| HFX | 39 | 0.788 | 0.212 | 100 | 100 | 100 | 97.4 |
| JRX | 58 | 0.704 | 0.296 | 100 | 100 | 100 | 84.5 |

Breeds with highest proportion of individuals with >6.25% taurine admixture included Punganur (44.4%), Vechur (42.3%), Umblachery (33.3%) and Pulikulam (23.5%) cattle. Significant proportion of individuals belonging to these breeds also had >12.5% taurine admixture. Among these, Vechur and Pulikulam have been reported to have severe shortage of breedable males (DADF, 2015; Singh and Sharma, 2017) in their respective native tracts. However, the AI coverage in these areas is high and the semen of purebred, commercial taurine cattle are readily available for insemination in local cows. This might have resulted in indiscriminate crossbreeding of purebred zebu cattle, thereby increasing the level of taurine admixture in them. In case of Punganur cattle, although reasonable number of breedable males are available in the native tract, the level of taurine admixture is relatively higher, possibly due to well established AI programs and readily available exotic taurine semen for insemination. Kannadhasan et al. (2018) utilized community based participatory approach to identify and prioritize the constraints in Umblachery cattle farming: (i) shift in farmers’ preference towards crossbreds for better milk production (ii) lack of interest among farmers in using Umblachery bullocks for agricultural operations (iii) lack of grazing lands (iv) lack of quality purebred breeding bulls and (iv) inadequate efforts of breeders’ association in promoting and improving Umblachery breed. The above factors coupled with high coverage of AI and easy access to taurine/crossbred semen might have contributed to relatively higher levels of taurine admixture in Umblachery cattle.

### 4.4. Maternal lineage diversity and zebu cattle domestication

The present study revealed no additional maternal haplogroups in the South Indian cattle other than the typical I1 and I2 zebu lineages reported earlier (Chen et al. 2010). A relatively high average number of nucleotide differences, nucleotide and haplotype diversity was observed in I2 lineage as compared to that of I1lineage. Mismatch distribution analysis of mtDNA control region sequences revealed the modal value at two mismatches for I1 lineage, while it was observed at two to four mismatches for I2 lineage in most of the South Indian breeds. The mismatch distribution analysis thus clearly showed high levels of mitochondrial control region diversity in South Indian zebu cattle. In general, extant populations located nearby centres of domestication retain high levels of diversity, while populations located farther away from the domestication centre show reduced diversity. The high levels of mtDNA control region diversity observed in South Indian zebu cattle is interesting. Particularly, the pairwise differences among the I2 haplotypes of South Indian cattle were relatively higher than West Indian (Indus valley site) zebu cattle and comparable to that of North Indian and Nepalese zebu cattle (Gangetic plains zebu) (Chen et al. 2010).

Several studies have demonstrated independent domestication and origin of *Bos tauus* and *Bos indicus* cattle based on mtDNA variations (Loftus et al. 1994). It is also clearly established that the humpless *Bos taurus* cattle were domesticated in the Near East during the Neolithic transition about 10 thousand years before present (Troy et al. 2001; Beja-Pereira et al. 2006; Edwards et al. 2007; Kantanen et al. 2009; Bonfiglio et al. 2010). Similarly, the report of Chen et al. (2010) shed much light on the origin and domestication of humped zebu cattle in the Indian sub-continent. They analyzed mitochondrial control region sequences from six different regions of Asia (i) Indus valley region (ii) Gangetic plains (iii) South India (iv) North East India (v) West Asia (vi) East Asia (vii) Central Asia and (viii) South East Asia. Among these, nucleotide diversity was reported to be notably higher in four regions of the Indian sub-continent (Indus Valley, Gangetic Plains, South India and North East India), indicating a vast expanse of area as the probable centre of domestication. Chen et al. (2010) identified the greater Indus valley (West India and present-day Pakistan) as most likely location for the origin and domestication of I1 maternal lineage. They made the above conclusion based on the following evidences: (i) the backbone structure of I1 haplogroup network being formed by cattle populations in the Indus valley region (ii) modal value of mismatch distribution of pairwise differences being higher in Indus valley cattle (two mismatches) as compared to those located in Gangetic plains (one mismatch) and South India (one mismatch) and (iii) relatively high I1 haplotype diversity observed in Indus valley cattle as compared to cattle in Gangetic plains and South India. However, analysis of I2 haplogroup sequences showed more complex pattern of diversity among cattle located in these regions. Identical modal values of I2 mismatch and nearly identical haplotype diversity were observed in Indus valley and South Indian cattle while the frequency of I2a (an intermediate I1-I2 haplotype) was relatively higher in cattle from Gangetic plains. Hence, Chen et al. (2010) reported the lack of genetic evidence to pin-point a single location/region for the origin of the I2 haplogroup. However, they opined the possible initial domestication of this lineage in the northern part of the Indian sub-continent based on the chronology of available archaeological evidence for domesticated zebu cattle and probable wild aurochs in the Indian subcontinent.

The present study based on comprehensive sampling of diverse breeds of South Indian zebu cattle, showed the mismatch modal value of I1 haplogroup to be higher (two mismatch) than reported by Chen et al. (2010). More interestingly, the mismatch distribution analysis of I2 haplogroup sequences revealed modal values around two to four mismatches in most South Indian breeds. This was higher than the modal value observed for West Indian (Indus valley) cattle, but comparable to that of North Indian and Nepalese (Gangetic plains) cattle. The intermediate I1-I2 haplotype sequences (I2a) were also observed in certain South Indian cattle breeds like Pulikulam, Deoni and Kangayam. Further, it is worth to mention that the haplotype diversity within South Indian I2 lineage was higher in most South Indian cattle breeds. Principal components analysis of pairwise F_ST_ derived from frequency of I2 haplotypes showed the distinctness of Vechur, Kangayam, Punganur and Alambadi cattle, thus indicating the diversity of this lineage within South Indian cattle. The results showed the need for additional sampling of zebu breeds from North, East and North-East Indian cattle to explore and gather new evidence on the potential location for the origin and domestication of I2 lineage in the Indian sub-continent.

## Conclusions

The present study revealed moderate levels of genetic diversity existing in South Indian zebu cattle. It also established the genetic distinctness of Kangayam, Vechur and Punganur cattle from the rest of the South Indian zebu cattle. The influence of Hallikar breed was observed in the development of most Mysore type cattle breeds of South India with the exception of Kangayam. Analysis of mtDNA control region sequences revealed two major maternal haplogroups, I1 and I2, with the former being predominant than the later. The results indicated the need for additional sampling and comprehensive analysis of mtDNA control region variations to unravel the probable location of origin and domestication of I2 zebu lineage. The present study revealed two major concerns about the status, conservation and improvement of South Indian draught type zebu cattle breeds: (i) Four of the ten investigated zebu cattle breeds are under the risk of endangerment due to small effective population size and high rate of inbreeding (ii) the lack of sufficient purebred zebu bulls for breeding and increasing level of taurine admixture in the native breeds tracts. Recently, the state governments of South India have started establishing conservation units and research centres for improvement of breeds like Alambadi, Bargur, Kangayam and Umblachery. Parental stocks of these breeds are being established by procuring true to the type animals from small holder farmers. However, in the absence of pedigree records, it has been difficult to estimate genetic purity or taurine admixture in those animals. Considering the relatively higher levels of taurine admixture in some of the breeds, it is advisable to perform molecular/genomic evaluation of local cattle to improve breed purity in these units/centres. It also needs to be mentioned that additional efforts are necessary to make the farmers aware of scientific breeding practices and limit the levels of inbreeding and taurine admixture in purebred zebu populations. Availability of purebred semen for artificial insemination, incorporation of genomic/molecular information to identify purebred animals and increased awareness among farmers will help to maintain breed purity, conserve and improve these important draught cattle germplasms of South India

## Acknowledgement

The present study is part of the Coordinated Research Project D3.10.28 of Joint FAO/IAEA Division of Nuclear Techniques in Food and Agriculture, International Atomic Energy Agency (IAEA), Vienna, Austria. The funding provided by IAEA for internship training of the first author at Animal Production and Health Laboratory, Seibersdorf, Austria is gratefully acknowledged. The authors also thank Dean, Veterinary College and Research Institute, Namakkal, Tamil Nadu, India for providing necessary facilities for the study.

## Legends of Supplementary Information

### Supplementary Figures

Supplementary Figure SF1. The test for selective neutrality at 27 microsatellite marker loci. Selection detection based on the *F*_ST_ outlier approach using LOSITAN showed that two marker loci (HEL13 and HEL5) deviated from selectively neutrality

Supplementary Figure SF2. Determination of correct number of clusters in Bayesian STRUCTURE analysis (Evanno et al. 2005) (a) Mean L (K) over 10 runs for each K value of 1 to 15 (b) Distribution of ΔK with the modal value (K=2) indicating the true K or the uppermost level of structure

Supplementary Figure SF3. Maximum likelihood tree of mitochondrial DNA haplotypes of draught type zebu, taurine and crossbred cattle from South India

### Supplementary Tables

Supplementary Table ST1. Details of 27 FAO recommended microsatellite loci used to evaluate South Indian draught cattle breeds

Supplementary Table ST2. Accession numbers, population information and haplogroup details of GenBank sequences used in the study

Supplementary Table ST3. Global F statistics among South Indian cattle (a) draught type zebu, taurine and crossbred cattle and (b) draught type zebu cattle only

Supplementary Table ST4. Frequency of mtDNA haplotypes and their sharing among different breeds of zebu, taurine and crossbred cattle

